# Structural basis of K11/K48-branched ubiquitin chain recognition by the human 26S proteasome

**DOI:** 10.1101/2025.01.13.632666

**Authors:** Piotr Draczkowski, Szu-Ni Chen, Ting Chen, Yong-Sheng Wang, Jessica Y. C. Huang, Ming-Chieh Tsai, Shu-Yu Lin, Rosa Viner, Yuan-Chih Chang, Kuen-Phon Wu, Shang-Te Danny Hsu

## Abstract

Beyond the canonical K48-linked homotypic polyubiquitination for proteasome-targeted proteolysis, K11/K48-branched ubiquitin (Ub) chains are involved in fast-tracking protein turnover during cell cycle progression and proteotoxic stress. Here, we report cryo-EM structures of human 26S proteasome in a complex with a K11/K48-branched Ub chain. The structures revealed a multivalent substrate recognition mechanism involving a hitherto unknown K11-linked Ub binding site at the groove formed by RPN2 and RPN10 in addition to the canonical K48-linkage binding site formed by RPN10 and RPT4/5 coiled-coil. Additionally, RPN2 recognizes an alternating K11-K48 linkage through a conserved motif similar to the K48-specific T1 binding site of RPN1. The insights gleaned from these structures explain the molecular mechanism underlying the recognition of the K11/K48 branched Ub as a priority signal in the ubiquitin-mediated proteasomal degradation.

## Introduction

Ubiquitin (Ub) is implicated in a myriad of biological processes owing to its ability to form multiple types of polymeric conjugates through one (or more) of the seven lysine (K) residues or the N-terminal methionine (M1). In addition to the well-studied homotypic Ub chains, atypical Ub chain topologies have been identified. These include heterotypic mixed-linkage Ub chains, encompassing more than one type of isopeptide bond linkage along a linear chain, and complex heterotypic branched Ub chains, encompassing Ub molecules in which more than one lysine is involved in the chain formation resulting in multiple branching points within the same Ub chain^1^. Branched Ub chains account for 10–20% of Ub polymers^2,3^. Among the different types of branched Ub chains, K11/K48-linked Ub branching is the best characterized: it is preferentially recognized by the ubiquitin-proteasome system (UPS) for substrate degradation under specific cellular conditions, including cell cycle progression in early mitosis and proteotoxic stress that requires proteostasis maintenance^4^. The K11/K48-branched Ub chains mediate the timely degradation of mitotic regulators, misfolded nascent polypeptides, and pathological Huntingtin variants^4–6^. The K11/K48-branched Ub chains adopt different topologies in a cellular context-dependent manner^4^. However, the mechanism by which the proteasomal Ub receptors recognize the K11/K48-branched Ub chain and differentiate it from its homotypic counterparts remains unknown.

The 26S proteasome recognizes ubiquitinated substrates through three constitutive Ub/Ub-like (UBL) receptors – RPN1, RPN10, and RPN13 – located within the 19S regulatory particle (RP) subcomplex^7–11^. RPN10 binds to Ub through two α-helical ubiquitin interacting motifs (UIMs) tethered to the N-terminal Willebrand factor A (VWA) domain within the 19S RP^7,12^. RPN13 binds to Ub through its N-terminal pleckstrin receptor for ubiquitin (PRU) domain tethered to the flexible C-terminus of RPN2 while the C-terminal part of RPN13 forms a deubiquitinase adaptor (DEUBAD) domain to recruit ubiquitin C-terminal hydrolase L5 (UCHL5; also known as UCH37)^8,9,13–16^. The PRU and DEUBAD domains of RPN13 are connected by a disordered linker, providing accessibility to a large conformational space for substrate recognition^17^. RPN1 binds to Ub at the T1 site, formed by a three-helix bundle within its proteasome/cyclosome (PC) domain^10^ or at its N-terminal section^11^. The enhanced binding of K11/K48-branched Ub chains was reported for isolated RPN1 and RPN10, which could account for the accelerated proteasomal degradation of substrates marked with K11/K48 Ub chains^6,18^. Additionally, Ub binding of all three proteasomal Ub receptors involves auxiliary factors such as shuttling factors and deubiquitinating enzymes (DUBs) that further contribute to Ub chain specificity^10,14,16,19,20^. In particular, the RPN13-associated UCHL5 preferentially recognizes and removes K11/K48-branched Ub chains from proteasomal substrates^21,22^.

In addition to the canonical proteasomal Ub receptors, there is good biochemical and genetic evidence to suggest the presence of one or more hitherto unidentified cryptic Ub receptors within the 26S proteasome^10,23^. While Ub binding to RPT5 and Sem1/Dss1 may account for the cryptic Ub receptor, how they bind to Ub remains poorly understood^24,25^. Recently, two potential Ub binding sites have been suggested: the α5-subunit within the 20S core particle (CP) subcomplex and RPN2 within the 19S RP. The latter is a paralog of the RPN1 that is structurally homologous to the T1 Ub binding site of RPN1^26^. UBL binding to RPN2 was observed by chemical cross-linking, but the molecular details remain elusive^27^. Importantly, whether the cryptic proteasomal Ub binding site is also implicated in recognizing branched Ub chains remains unknown.

Despite the wealth of biochemical and structural understanding of the UPS, chiefly deduced from recent cryo-EM studies^28–40^, the molecular basis of how branched Ub chains are recognized and processed by the 26S proteasome remains poorly understood. Here, we report the cryo-EM structures of the human 26S proteasome bound to a substrate conjugated with a K11/K48-branched Ub chain. We unambiguously resolved the cryo-EM structure of a tetra-ubiquitin with a K11/K48 branching point at the proximal Ub to form a well-defined tripartite binding interface with the 19S RP. The cryo-EM structures provided direct structural evidence to indicate RPN2 as a Ub receptor recognizing the K48-linkage extending from the K11-linked Ub, forming a unique alternating K11-K48 linkage stemming from the proximal Ub that helps position the K11-linked Ub branch into a groove formed by RPN2 and neighboring proteasomal subunits. The new structural insights from this study illustrate the versatility of the 26S proteasome in decoding the complex Ub chain signaling needed to maintain proteostasis.

## Results

### Biochemical and structural characterization of a K11/K48-branched ubiquitin chain bound to the 26S proteasome complex

We reconstituted a functional complex of the human 26S proteasome in a complex with a polyubiquitinated substrate and the auxiliary proteins RPN13 and UCHL5. The substrate consisted of the intrinsically disordered residues 1-48 of *S. cerevisiae* Sic1 protein (hereafter Sic1^PY^)^39^ with a single lysine residue (K40) that serves as an anchoring point for ubiquitination by an engineered Rsp5 E3 ligase (hereafter Rsp5-HECT_GML_; Materials and Methods; Fig. 1a). While wild-type (WT) Rsp5 generates K63-linked Ub chains^41^, Rsp5-HECT_GML_ has been engineered to generate K48-linked Ub chains, which was confirmed by Western blotting using Ub linkage-specific antibodies^42^. To exclude the possibility of generating a K63-linked Ub chain, a Ub variant (K63R) of which the lysine at position 63 is mutated into an arginine was used to produce the polyubiquitinated Sic1^PY^ (Sic1^PY^-Ub_n_) (Supplementary Fig. 1). Dual fluorescence labelling for Sic1^PY^ (Alexa488) and Ub (fluorescein) was introduced to enable simultaneous detection of the two, which helps distinguish substrate proteolysis from deubiquitination. Sic1^PY^-Ub_n_ stably bound to the enzymatically active human 26S proteasome (Supplementary Fig. 2). The crude Sic1^PY^-Ub_n_ reaction product was further fractionated by size-exclusion chromatography (SEC) to enrich the medium-length Ub chains (n = 4-8) to be efficiently processed by the 26S proteasome (Fig. 1b and Supplementary Fig. 2). Contrary to the expectation of forming predominantly K48-linked homotypic Ub chains, however, Lb^pro*^ Ub clipping^2^ and intact mass spectrometry (MS) analysis revealed the presence of doubly ubiquitinated (12.6%) and triply ubiquitinated (3.6%) Ub in addition to singly ubiquitinated Ub (41.8%), which is clear evidence of the formation of branched Ub chains (Fig. 1c). To identify the linkage types of the poly-Ub chains, we employed MS-based Ub absolute quantification (Ub-AQUA)^43,44^ to demonstrate that the SEC-enriched poly-Ub chains contained almost equal amounts of K11- and K48-linked Ub with a minor population of K33-linked Ub (Fig. 1c and Supplementary Fig. 3d). By contrast, there was less K11-linked Ub before the SEC enrichment (Supplementary Fig. 3b). Additionally, intact MS analysis of the Lb^pro*^ trimmed poly-Ub chains before SEC enrichment showed marginally decreased amounts of doubly ubiquitinated (11.6%) and triply ubiquitinated (1.8%) Ubs compared to SEC-enriched short Ub-chains (Supplementary Fig. 3a and 3c).

**Fig. 1.**
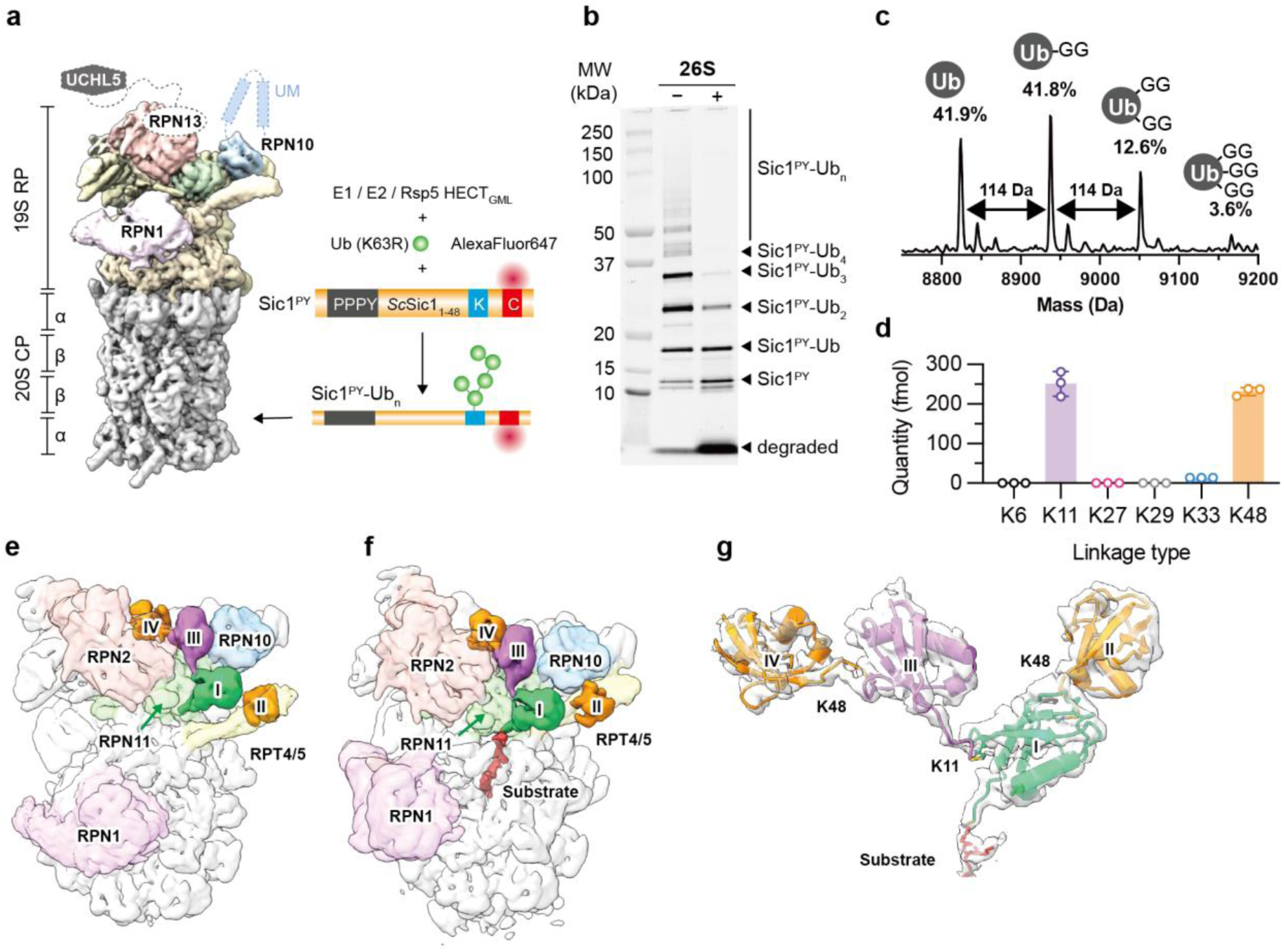
Reconstitution and cryo-EM analysis of the human 26S proteasome in complex with a K11/K48-branched Ub chain-modified Sic1^PY^. **a**, Construct design and labeling strategy of a polyubiquitinated Sic1^PY^. **b**, 26S proteasome-mediated proteolysis of the polyubiquitinated Sic1^PY^ detected by SDS-PAGE separation followed by fluorescence imaging of the exogenously labeled AlexaFluor647. **c**, Evidence of Ub chain branching by Lb^pro*^ digestion followed by intact MS analysis. The difference of 114 Da is indicative of a single ubiquitination site on the target Ub. The relative populations in percentage are derived from peak integration of the individual peaks. The experiments were carried in triplicate with the original data points shown in open circles and the error bars indicating the standard deviations. **d**, MS Ub-AQUA/PRM quantification of the abundances of the individual Ub chain linkage types. K63 is omitted due to the use of the Ub(K63R) variant. The experiments were carried in triplicate with the original data points shown in open circles and the error bars indicating the standard deviations. Cryo-EM maps of the reconstituted proteasomal complex in the E_B_ state (**e**), and substrate-processing E_D_ state (**f**), Well-resolved cryo-EM map of the K11/K48-branched Ub_4_ chain can be identified in both states. **g**, Segmented cryo-EM map of the proteasome-bound Ub_4_ in the ED state is superimposed with the atomic model to illustrate the clear definition of the K11/K48-branched Ub chain topology. The positions of the Ub_I_, Ub_II_, Ub_III_ and Ub_IV_ are indicated in Roman numbers.

UCHL5 is a proteasome-associated DUB that preferentially processes K11/K48-branched Ub chains^21^. The DUB activity of UCHL5 is activated upon binding to RPN13^8^. The addition of UCHL5 to the reconstitute proteasomal complex may help capture K11/K48-branched Ub chains. However, UCHL5 could also disassemble the proteasome-bound Ub chains. To minimize the processing of the Sic1^PY^-Ub_n_ by the endogenous UCHL5, we added an excess amount of preformed RPN13:UCHL5 complex with an alanine mutation for the catalytic cysteine of UCHL5(C88A). The presence of UCHL5, RPN13 and Sic1^PY^-Ub_n_ in the reconstituted complex was confirmed by native gel electrophoresis combined with Western blotting and fluorescence imaging (Supplementary Fig. 2). The ternary functional complex formation was additionally confirmed by negative staining electron microscopy (NSEM), which showed additional EM densities on the 19S RP of the reconstituted proteasomal complex when compared to the apo 26S proteasome (Supplementary Fig. 2d).

We successfully determined four cryo-EM structures of the reconstituted proteasomal complex after extensive classification and focused refinements (Fig. 1d-f, Supplementary Fig. 4 and 5, and Supplementary Movie 1). These structures resembled the previously reported substrate-free (apo) E_A_ state, an Ub chain-bound E_A_, E_B_, and substrate-engaged E_D_ state of human proteasome^37^. Importantly, we unambiguously resolved the cryo-EM densities of a K11/K48-branched tetra-Ub (Ub_4_) chain bound to the E_B_ and E_D_ states (hereafter E_B_:Ub_4_ and E_D_:Ub_4_, respectively;). For the Ub chain-bound E_A_ state, the most distal Ub could not be resolved in the cryo-EM map, but the core K11/K48-branched tri-Ub remained well-defined (hereafter E_A_:Ub_3_). Additionally, the E_D_:Ub_4_ state exhibited extra density extending from the proximal Ub into the base of the 19S RP., corresponding to the Sic^PY^ substrate poised for *en bloc* deubiquitination before being translocated into the 20S core particle (CP) for degradation (Fig. 1e).

Despite the addition of an excess amount of UCHL5(C88A), we did not observe EM density that could be assigned to RPN13-bound UCHL5 in any of the four functional states. This may be due to the intrinsic dynamics of RPN13 and UCHL5 on the proteasome^17,45,46^. Indeed, in the absence Sic1^PY^-Ub_n_, we identified by chemical cross-linking MS analysis several contacts between UCHL5 and proteasomal proteins that are spatially distant from each other (Supplementary Fig. 6). In the presence of Sic1^PY^-Ub_n_, the chemical cross-linking of UCHL5 was limited to Ub, suggesting the displacement of UCHL5 by Sic1^PY^-Ub_n_ (Supplementary Fig. 6). The intrinsic dynamics of UCHL5, RPN10 UIM motifs (Fig. 1a), and RPN13 potentially allow the complex to target a wide range of ubiquitinated substrates. Nonetheless, such functional dynamics also makes it intrinsically challenging to define their precise locations on the proteasome^47,48^.

### Structural basis of K11/K48-branched Ub chain bound to the 26S proteasome with multivalency

The proximal Ub in the E_A_:Ub_3_, E_B_:Ub_4_ and E_D_:Ub_4_ all adopted a conserved pose similar to several previously reported RPN11:Ub structures (Supplementary Fig. 7a). The K48 of the proximal Ub was linked to another Ub in contact with the RPT4/5 coiled-coil (CC) adjacent to RPN10 (Fig. 1e). The relative orientation of the RPT4/5 CC-bound K48-linked Ub was similar to previously reported cryo-EM densities of distal proteasome-bound Ub (Supplementary Fig. 7b)^37,38^, in line with the previously reported role of RPT5 in polyUb-binding by the proteasome^24^. Despite its dynamics, the density of the K48-linked Ub exhibited a clear Ub fold – a signature thumb and palm shape – enabling the modeling of the interactions between the K48-linked Ub and proteasomal subunits (Supplementary Fig. 7c).

In addition to the canonical K48-linked Ub, we identified an additional Ub linked to the proximal Ub through a K11 linkage to form a K11/K48-branched Ub chain topology (Fig. 1f). The K11-linked Ub nicely tucked into a multi-subunit groove encompassing RPN2, RPN8, RPN9, RPN10, and RPN11. The ability of this novel Ub binding groove to bind K11-linked Ub was further confirmed by a control cryo-EM analysis of which Sic1^PY^ was modified by a K11-only homotypic Ub chain. A well-resolved cryo-EM density was identified in the newly identified Ub binding groove but not in the RPT4/5 CC region (Supplementary Fig. 8). Compared to previously reported K11-linked di-Ub structures, proteasome-binding required substantial conformation rearrangements of the K11-linked Ub relative to the proximal Ub to dock into the newly identified Ub binding groove (Supplementary Fig. 7d)^47,49^. For simplicity, the proximal Ub, the K48-linked Ub, and the K11-linked Ub that formed the core K11/K48-branched Ub chain are hereafter defined as Ub_I_, Ub_II_, and Ub_III_, respectively (Fig. 1d-f)

In addition to the K11/K48-branched Ub_3_ core, we resolved the fourth Ub (defined as Ub_IV_).) in the E_B_:Ub_4_ and E_D_:Ub_4_ states. Ub_IV_ was linked to Ub_III_ through a K48-linkage, resulting in an alternating K11-K48 linkage stemming from Ub_I_ (Fig. 1d-f). To the best of our knowledge, such an alternating linkage has not been documented in the literature. Collectively, our cryo-EM analysis revealed a novel multivalent binding mode between the human 26S proteasome and the K11/K48-branched Ub chain encompassing the canonical K48-link Ub binding site and the newly identified binding site for the K11-linked Ub chain.

### Alternating K11-K48 linkage along the K11-linked branch of the Ub chain allows for an extensive interaction interface with the proteasomal 19S RP

Close examination of the molecular details of the interactions between the K11/K48-branched Ub chain and the proteasome revealed an extensive Ub interaction network involving RPN2, RPN8, RPN9, RPN10, RPN11, and the RPT4/5 CC in the E_A_:Ub_3,_ E_B_:Ub_4_ and E_D_:Ub_4_ states. The multi-subunit binding interface sequestered a bulk part of the solvent-accessible surface area (ΔSASA) of the K11/K48-linked Ub_4_ (Supplementary Table 1-3). The intermolecular hydrogen bonds involved C-terminal residues of Ub_I_, particularly in the E_A_:Ub_3_ and E_B_:Ub_4_ states (Fig. 2 and ^S^upplementary Table ^1^). This is in line with the general functional role of RPN11 in removing the Ub chains *en bloc* from target substrates^50^. In both the E_B_:Ub_4_ and E_D_:Ub_4_ states, the I44 and I36 hydrophobic patches of Ub_I_ made extensive contacts with the hydrophobic lining of the active site of RPN11 (Fig. 2). In contrast, Ub_II_ made limited contacts with RPN10 and the RPT4 RPT5 CC primarily through hydrogen bonding. Notably, between the E_B_:Ub_4_ and E_D_:Ub_4_ states, the RPT4/5 CC underwent a pronounced lever motion over which Ub_II_ hopped over to contact opposite sides of the RPT4/5 CC: RPT5 was involved in hydrogen bonding with Ub_II_ in the E_B_:Ub_4_ state while RPN10 was involved in hydrogen bonding with Ub_II_ in the E_D_:Ub_4_ state (Fig. 2 and Supplementary ^T^able ^1^).

**Fig. 2.**
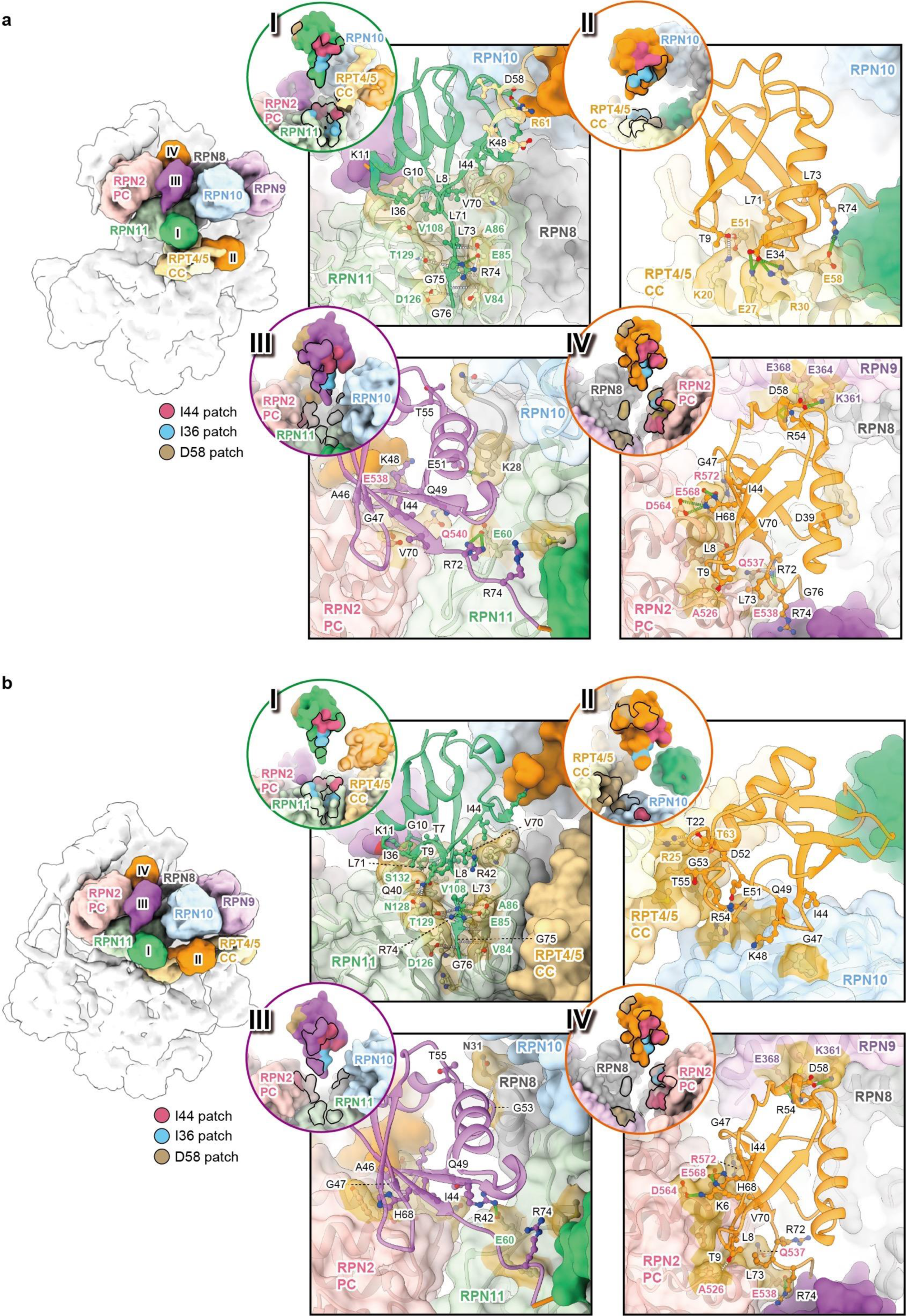
Structural basis of K11/K48-branched Ub chain recognition by the human 26S proteasome. Cryo-EM structures of the 19S RP in the E_B_:Ub_4_ state (**a**) and substrate-processing E_D_:Ub_4_ state (**b**) bound with the K11/K48-branched Ub_4_ with the same coloring scheme as in Fig. 1e. The expanded views of the individual Ubs in the two functional states are shown on the right panels with the individual Ub shown in cartoon representations and the proteasomal proteins in surface representations. The identities of the interacting residues in Ub are labeled in black. The identities of the proteasomal residues involved in forming salt bridges and/or hydrogen bonds with Ubs are indicated with matching colors according to the scheme shown on the left panels. Circled insets on the upper left corners of individual Ub panels illustrate the Ub binding interfaces outlined by black lines. The I44, I36 and D58 patches on the Ubs are colored hot pink, blue and brown, respectively. Proteasomal residues involved in contacting the I44, I36 and D58 patches are colored in the same way.

Regarding the newly identified K11-Ub chain binding groove, Ub_III_ formed limited hydrogen bonds with RPN2 and RPN11 in all three functional states; Ub_III_ made additional hydrogen bonds with RPN8 only in the E_D_:Ub_4_ state (Fig. 2 and Supplementary Table 1). In contrast to the sparse interactions formed by Ub_II_ and Ub_III_, Ub_IV_ made extensive contacts with RPN2 and RPN9 (Fig. 2a-b panel IV, Fig. 3, Supplementary Table 1 and 2). Among all the four Ubs, Ub_IV_ contributed to the second largest ΔSASA (after the proximal ubiquitin, Ub_I_), accounting for 28% and 25% of the overall ΔSASA occupied by the entire K11/K48-linked Ub_4_ in the E_B_:Ub_4_ and E_D_:Ub_4_ complexes, respectively (Supplementary Table 3). Given the sparse contacts made by the K11-linked Ub_III_ with RPN2, the primary role of Ub_III_ appeared to restrict the conformational space available to Ub_IV_ for positioning it to the new Ub binding groove formed by RPN2 and RPN9. As a result, the unique alternating K11-K48 Ub linkage provided two anchoring points – Ub_I_ and Ub_IV_ – with optimal conformational complementarity with the 26S binding sites (Fig. 2). Collectively, the K11/K48-branched Ub_4_ contributed to a large ΔSASA of 3077 and 2938 Å^2^ within the 19 RP of the E_B_:Ub_4_ and E_D_:Ub_4_ states, respectively (Supplementary Table 3).

**Fig. 3.**
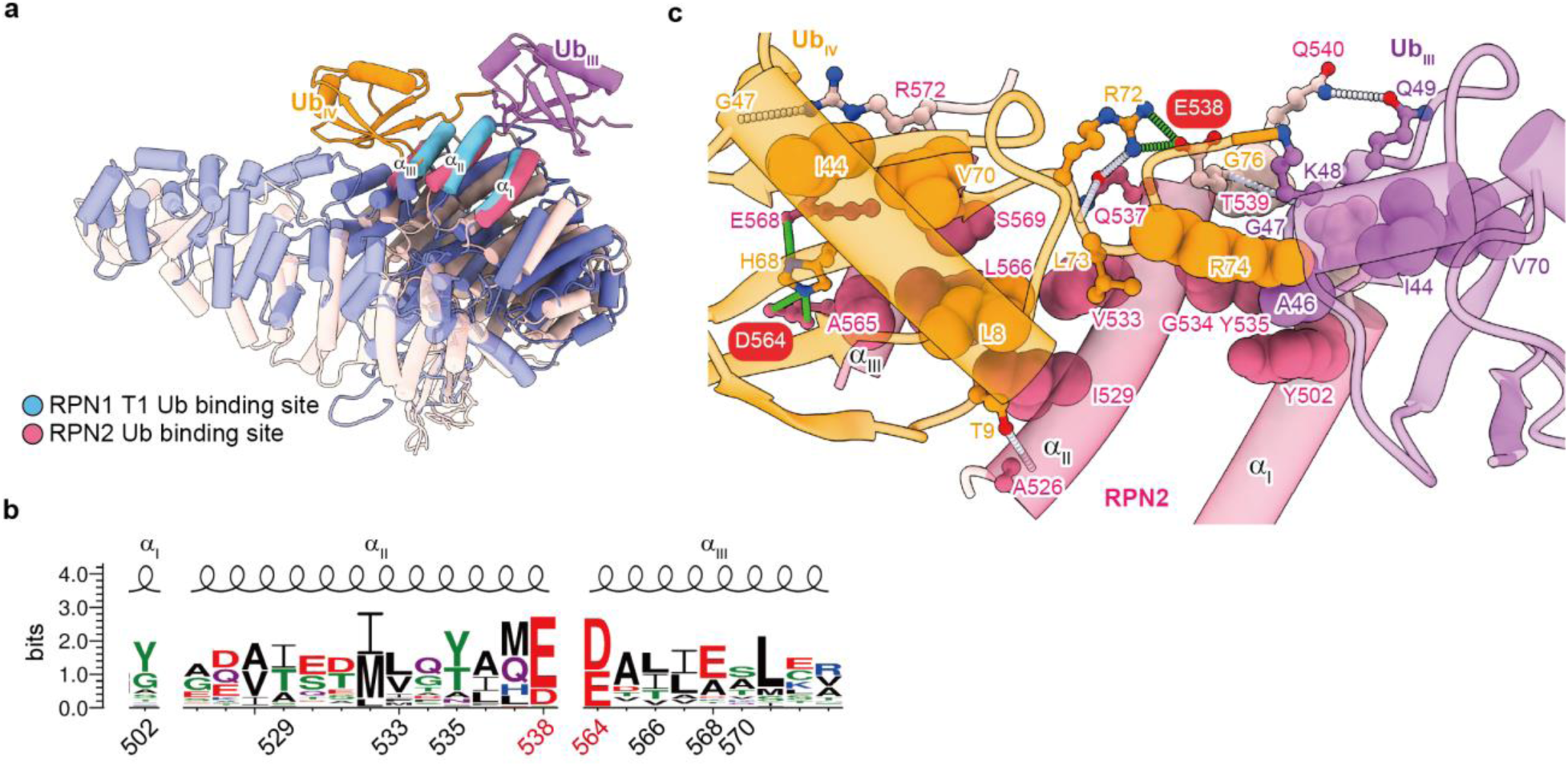
Structural basis of K48-linked di-Ub recognition by RPN2. **a**, Structural alignment of RPN1 (orchid) and RPN2 (fuchsia) focusing on the helical Ub binding motifs (highlighted in blue and hot pink, respectively). The alternating K11-K48-linked di-Ub is colored following the same scheme as in Fig. 1e. The structure of RPN1 is taken from the PDB entry 6J2N. The structure of the RPN2 in complex with the K11-K48-linked di-Ub is taken from the E_D_:Ub_4_ state in this study. **b**, Expanded view of the K48-linked di-Ub in complex with RPN2. The conserved E538 and D564 are highlighted by red ovals. The inter-chain hydrogen bonds and salt bridges are shown in dashed white lines and dashed green lines, respectively. c, LOGO representation of the sequence conservation of the K48-linkage binding motif shared by RPN2 and RPN1. The conserved acidic residues are indicated in red, and the other conserved hydrophobic residues are highlighted in black below the sequence alignment. They are numbered according to the sequence of RPN2.

### RPN2 contains a conserved ubiquitin-binding motif

Our cryo-EM structures underscored the pivotal role of RPN2 in recognizing the K11-linked Ub branch with alternating K11-K48 linkages, namely Ub_III_-Ub_IV_. Close examination of the structural motifs pertinent to Ub recognition revealed a conserved K48-linked di-Ub recognition motif within RPN2. The motif consists of three α-helices within the PC domain of RPN2, resembling the canonical T1 Ub receptor binding motif of RPN1^10^ (Fig. 3a). Alignment of 23 RPN1 and 21 RPN2 sequences from different model organisms revealed two highly conserved negatively charged residues involved in Ub binding, corresponding to E538 and D564 in human RPN2 (Fig. 3c and Supplementary Fig. 9).

In the E_B_:Ub_4_ state, E538 of RPN2 formed a salt bridge with R72 of Ub_IV_, while D564 formed a salt bridge with H68 of Ub_IV_ (Fig. 3b). The interaction was consolidated by R572 whose side-chain was hydrogen bonded to the backbone carbonyl of G47; concomitantly, Q537 made contacts with R72 and L73 of Ub_IV_ (Fig. 3b). The conserved hydrophobic surface encompassing A526, I529, V533, G534, Y535, A565, L566, and S569 made extensive contacts with L8, T9, I44, V70, L73, and R74 of Ub_IV_ (Fig. 3b). In contrast, in the E_D_:Ub_4_ state, the interaction network between the RPN2 and Ub_IV_ underwent rearrangements resulting in reduced intermolecular hydrogen bonds and salt bridges (Supplementary Table 1). Specifically, E538 of RPN2 formed a salt bridge with R74 of Ub_IV_, while D564 of RPN2 switched its salt bridge register to K6 of Ub_IV_. An additional salt bridge was formed between H68 of Ub_IV_ and the conserved E568 of RPN2 in the E_D_:Ub_4_ complex, while the Q537 of RPN2 switched its hydrogen bonds to the backbone amide group of L73 instead of R72 in the E_B_:Ub_4_ state. The hydrophobic interactions between RPN2 and Ub_IV_ were generally conserved between the E_B_:Ub_4_ and E_D_:Ub_4_ states, with the latter exhibiting an additional contribution from the conserved L570 of RPN2.

In contrast to the extensive interactions between RPN2 and Ub_IV_, RPN2 only made few interactions with Ub_III_. These included a hydrogen bond between Q540 of RPN2 and Q49 of Ub_III_, which existed in the E_B_:Ub_4_ state but not in E_D_:Ub_4_ state. Additionally, the hydrophobic interactions were formed mostly between the T539 of RPN2 and I44, A46, G47, and K48 of Ub_III_. Overall, the amphipathic nature of the di-Ub binding mode observed in RPN2 was similar but not identical to the one used by RPN1 (Supplementary Fig. 10). Importantly, this novel di-Ub binding motif of RPN2 could account for the previously proposed cryptic Ub receptor on the 19S RP in addition to RPN1, RPN10, and RPN13.

### Substrate binding triggers a cascade of conformational changes in the 19S RP

The 19S RP exists in many functional states with distinct conformational arrangements of the individual subunits. Our cryo-EM data revealed four distinct functional states that resemble the previously reported conformational states of the human 26S proteasome, namely the E_A_, E_B_, and E_D_ states^37^ (Supplementary Fig. 11). These functional states bound to Ub_3_ or Ub_4_ to form the E_A_:Ub_3_, E_B_:Ub_4_, and E_D_:Ub_4_ state (Fig. 4, Supplementary Fig. 5 and Supplementary Fig. 12). The initial substrate binding to the apo E_A_ state did not induce discernable conformational changes: the overall conformation of the 19S RP did not change from the apo E_A_ state to the E_A_:Ub_3_ state. The E_B_:Ub_4_ state underwent subtle but significant conformational changes relative to the E_A_:Ub_3_ state, especially for RPN1, RPN2 and the RPT4/5 CC (Fig. 4a). These subunit motions were accompanied by an overall tilting motion of the 19S RP lid, resulting in a pronounced displacement of the centre of mass (COM) of RPN2 by 11 Å (Supplementary Fig. 12a). Such a motion opened up the space between the 19S lid and the oligosaccharide/oligonucleotide binding (OB) ring of the ATPase RPT subunits, thereby facilitating substrate access to the AAA+ ATPase entry pore.

**Fig. 4.**
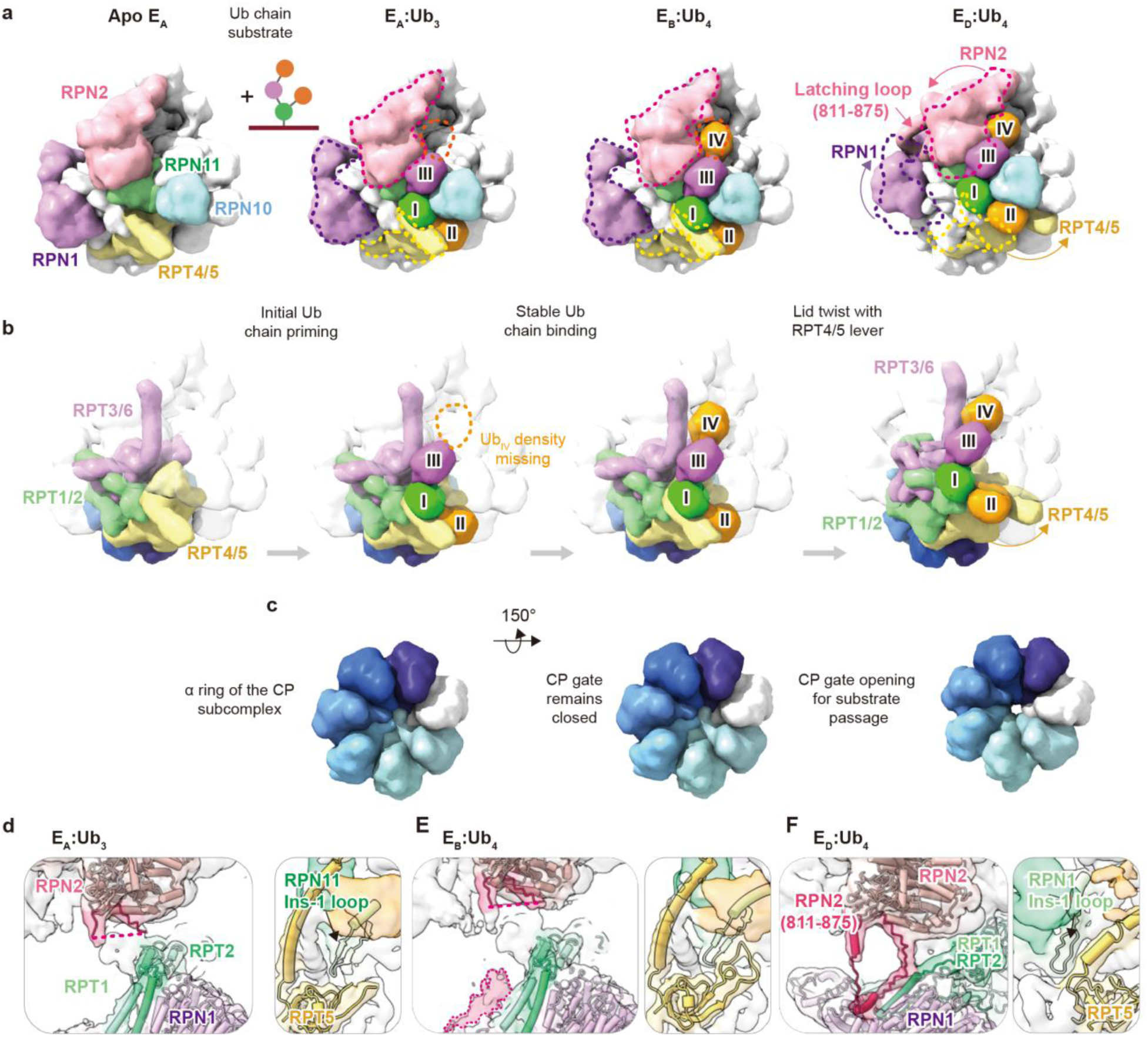
Conformational transitions of the 19S RP induced by substrate binding. **a**, Structural comparison of the 19S RP in the apo E_A_ state, E_A_:Ub_3_ state, E_B_:Ub_4_ state, and E_D_:Ub_4_ state following the same rendering scheme as in Fig. 1. The structures of the individual domains are purposely low-pass filtered to a resolution of 15 Å by ChimeraX according to their atomic models to highlight the large domain motions and CP gate alignment without the interferences of the finer details. For the Ub-bound states, the conformations of RPN1, RPN2 and RPT4/5 in the preceding states are outlined in dashed lines to highlight the rigid-body motions associated with the functional state transitions. The missing Ub_IV_ in the E_A_:Ub_3_ state is indicated by an orange dashed line. **b**, All 19S RP subunits but RPT1/2, RPT3/6 and RPT4/5 are shown in transparent surfaces to highlight the lever motions of the individual CCs, especially for the transition of the RPT4/5 CC from the E_B_:Ub_4_ state to the E_D_:Ub_4_ state. **c**, CP gate opening for substrate passage into the proteolytic chamber in the E_D_:Ub_4_ state. The seven subunits of the α-ring are color-ramped from white to dark indigo. Expanded views of the latching loop of RPN2 (residues 811-875; left panel) and the RPN11 Ins-1 loop (residues 76-88; right panel) in the E_A_:Ub_3_ state (**d**), E_B_:Ub_4_ state (**e**), and E_D_:Ub_4_ state (**f**). These intrinsically disordered loops are sufficiently rigidified and visible to cryo-EM analysis only in the E_D_:Ub_4_ state.

Finally, the E_D_:Ub_4_ state underwent the most substantial conformational changes in the entire 19S RP lid, with the most notable being the lever motion of the RPT4/5 CC leading to a switch of Ub_II_ binding mode (Fig. 4b). Furthermore, the CP gate opened up in this state upon the appropriate alignment of the AAA+ ring, the OB ring, and the α-subunit ring of the 20S CP, thereby allowing substrate passage into the proteolytic chamber for degradation (Fig. 4c and Supplementary Fig. 12b), which is the hallmark of the E_D_ state^37^.

In addition to the rigid-body motions of the individual subunits, the E_B_:Ub_4_ state exhibited partially resolved EM density of the intrinsically disordered residues 811-875 of RPN2 as a result of its interaction with the RPT1/2 CC (Supplementary Fig. 10c). This loop became fully resolved in the E_D_:Ub_4_ state. The structure of the E_D_:Ub_4_ state illustrated how the intrinsically disordered loop latched onto RPN1 near the RPT1/2 CC to help maintain the inter-subunit geometrical arrangements (Fig. 4d-f and Supplementary Fig. 12c). Additionally, the Ins-1 loop of RPT11, pertinent to the activation of DUB activity of RPN11^51^, also became resolved when transiting from the E_A_:Ub_3_ state to the E_D_:Ub_4_ state, facilitated by the loop hairpin (residues 54-74) of RPT5 in the E_B_:Ub_4_ state (Fig. 4d-f and Supplementary Fig. 12d). Collectively, the structural insights gleaned from the four distinct functional states underscored the dynamic interplay and cross-talk between the individual subunits within the 19S PR and 20S CP.

## Discussion

In this work, we have generated a K11/K48 branched Ub chain-modified Sic1^PY^ using an engineered chimeric Rsp5 Ub E3 ligase, Rsp5-HECT_GML_^42^. The substrate could be efficiently recognized and processed by the human 26S proteasome (Fig. 1 and Supplementary Fig. 2). We unequivocally determined the cryo-EM structures of the K11/K48 branched Ub chains with the branching point located at the proximal Ub, based on which we revealed how the K48-linked Ub_IV_ extended from the K11-linked Ub_III_ to form a hitherto unseen alternating K11-K48 linked Ub chain stemming from the proximal Ub_I_ (Fig. 1e-g). RPN2, RPN8, RPN9, RPN10, and RPN11 collectively form a multi-subunit groove to accommodate the K11-linked Ub chain. The K11-linkage specificity of this multi-subunit binding groove was corroborated by the cryo-EM analysis of the K11-only homotypic Ub chain (Supplementary Fig. 8). The K11/K48-branched Ub chain made extensive hydrogen bonds and hydrophobic interactions with the 19S RP through the canonical I36 and I44 hydrophobic patches within the individual Ubs (Fig. 2). Specifically, the K11-K48 linked di-Ub was recognized by conserved Ub binding motif within RPN2, reminiscent of T1 Ub binding site on RPN1 (Fig. 3). Unlike RPN1, which binds to K48-linked di-Ub using the T1 helical bundle alone^10^, the recognition of the K11-linked Ub chain involves several proteasomal subunits in addition to RPN2: Ub_III_ formed several salt bridges with RPN10 and RPN11; Ub_IV_ formed several salt bridges with RPN9 (Fig. 2). RPN2 alone may not be sufficient to bind to the alternating K11-K48-linked Ub chain stably. In other words, the multivalent nature of this unique Ub chain binding groove may explain why previous studies have failed to identify this cryptic Ub receptor because this binding groove consists of a collection of proteasomal subunits^10,23^.

Despite the intrinsic heterogeneity of the Ub chain length and linkages of the Sic1^PY^-Ub_n_ used in this study, the ability to resolve the cryo-EM maps of the branched Ub_4_ is a testament that the unique multivalent binding mode provides superior avidity over homotypic Ub chains and other forms of Ub linkages, facilitating *in silico* enrichment of particle images to yield the well-defined cryo-EM maps of the K11/K48-branched Ub chains (Supplementary Fig. 4 and 5). Superposition of previously reported Ub chain structures extending from the structure of Ub_IV_ showed steric clashes between the extended Ub moieties and RPN9 that line the bottom of the K11-Ub binding groove (Supplementary Fig. 13). It is plausible that UCHL5, known to edit K11/K48-branched Ub chains^21^, may function as a molecular ruler to trim additional Ub moieties extending from Ub_IV_ to avoid these steric clashes. However, despite the addition of the catalytically inactive UCHL5, we could not map its position on the substrate-bound complex. Indeed, Walkers and co-workers concluded in their recent cryo-EM study that the Ub binding sites on RPN10 and RPN13, which tethers the UCHL5 to the proteasome, are too dynamic to be underpinned by cryo-EM^38^.

The structural snapshots of the free and substrate-bound 26S proteasome revealed abundant and long-range conformational changes associated with the Ub-receptor binding and the catalytic cycle of the 26S proteasome. Between the E_B_:Ub_4_ state and E_D_:Ub_4_ state, the RPT4/5 CC underwent a significant lever motion to expose Ub_II_. The lever motion was accompanied by the CP gate opening accomplished by the appropriate alignment of the basal ring and the OB ring (Fig. 4c and Supplementary Fig. 12). The conformational changes within the ATPase assembly were reminiscent of the previously reported activation mechanism of archaeal PAN ATPase^52^, and the activation mechanism of the ATPase unfoldase activity upon Ub binding to the proteasomal Ub receptors^53^. The extensive tripartite binding interface formed by the K11/K48-branched Ub chain may generate a favorable driving force to trigger these allosterically coupled conformational changes together with the ordering of several intrinsically disordered loops (Fig. 4 and Supplementary Fig. 12). Our structural findings thus suggested a pivotal role of the ATPase base of the 19S RP as a hub to sense and transmit allosteric regulatory signals triggered by substrate binding^31,35,54,55^.

In summary, our study provides insight into how the human 26S proteasome recognizes a K11/K48-branched Ub chain through multivalent interactions involving a large number of proteasomal subunits, namely RPN2, RPN8, RPN9, RPN10, RPN11 and the RPT4/5 CC. The newly identified K11-Ub binding groove on the 19S RP is particularly complex. While we have identified a conserved K48-linked Ub binding motif on RPN2, which is structurally similar to that of the T1 motif of RPN1, RPN2 alone is not sufficient to form a stable complex with the unique alternating K11-K48-linked Ub chain. Additional contacts made by RPN8, RPN9, and RPN10 may be necessary to generate sufficient multivalency (Fig. 2, Fig. 3a, and Supplementary Fig. 9 and 10). This multi-subunit Ub binding groove helps account for the previously speculated cryptic Ub receptor^10,23^. Collectively, the K11/K48-branched Ub chain forms a large interaction surface with the proteasome that affords avidity, providing expedited turnover of K11/K48-branched Ub conjugated substrates at specific biological checkpoints for maintaining proteostasis in a timely fashion^4,6^.

## Materials and Methods

### Plasmids, genes, and site-directed mutagenesis

The plasmids pRSFduet_His-TEV-gsggs-Ub(WT), pRSFduet_His-TEV-gsggs-Ub(K0), pET21d_His-hE1, pETduet_His-SUMO-gg-UBCH7, pGEX_TEV-SENP2, pMBP_TEV-Rsp5-WW3-HECT, and pMBP_TEV-Rsp5 WW3-HECT_GML were synthesized by (GeneScript, USA). pET19b_RPN13 was a gift from Joan Conaway & Ronald Conaway (Addgene plasmid #19423; http://n2t.net/addgene:19423; RRID: Addgene_19423)^16^. pOPINB-AMSH* was a gift from David Komander (Addgene plasmid #66712; http://n2t.net/addgene:66712; RRID: Addgene_66712)^56^. Plasmid encoding the N-terminal hexahistidine (His_6_)-UCHL5(WT) gene was taken from the in-house plasmid collection. Plasmids encoding His_6_-TEV-gsggs-Ub(K63R), - Ub(K48R), -Ub(K11-only), -Ub(K48-only) and His6-UCHL5(C88A) were constructed using site-directed mutagenesis. K11-only or K48-only referred to the Ub variants of which all lysines but K11 or K48 were mutated into arginines, respectively. For each of the mutations, a pair of complementary mutagenic primers were designed with which the entire plasmid was amplified in a thermocycling reaction using a KOD (Invitrogen, USA) high-fidelity DNA polymerase and the plasmids encoding His_6_-TEV-gsggs-Ub(WT), -Ub(K0), or His_6_-UCHL5(WT) respectively as a template. The synthetic gene encoding the substrate polypeptide (later called Sic1^PY^) exhibiting an efficient degradation initiation region for the proteasome engagement was derived from the C-terminal sequence of *S. cerevisiae* Sic1 protein (residues 1-48). The sequence was modified to contain an additional PPxY motif and single lysine for ubiquitination by the Rsp5-E3 Ub ligase, a single cysteine for site-specific labeling, and His_6_-tag at its C-terminus. The gene cloned in pET-9a plasmid by *Nde*I/*Bam*HI was ordered from GenScript, USA.

### Proteins and protein purification

All recombinant proteins used in the study, except human His_6_-RPN13, were expressed in *E. coli* BL21-CodonPlus (DE3)-RIL competent cells (Agilent Technologies). Proteins were expressed and purified following the protocol described below unless otherwise stated. During the purification procedures, protein solutions were kept on ice where possible, except for size-exclusion chromatography (SEC), during which the columns were kept at room temperature. A HiLoad 16/600 Superdex 200 pg column (GE Healthcare, USA) connected to an FPLC system (ÄKTA Purifier UPC 10 or ÄKTA Pure 25L, GE Healthcare, USA) was used for SEC unless otherwise specified. Human ubiquitin-activating enzyme His_6_-Uba1 (E1) was purified at 4 ℃ at all times, including the SEC steps.

After harvesting by centrifugation at 4 ℃, 6,000 g for 15 min using a JLA 8.1 rotor in Avanti −26 XP Series centrifuges. (Beckman Coulter, USA), cells were lysed using NanoLyzer N2 homogenizer (Gogene Corporation, Taiwan) operating at 18-20 kpsi, unless otherwise stated. After lysis, cell debris, and aggregates were removed by 30 min centrifugation at 4 ℃, 20,000 g using a JA 25.5 rotor in Avanti −26 XP Series centrifuges. (Beckman Coulter, USA). All the His-tag-based affinity purifications were done using the Ni-NTA resin (cOmplete™ His-Tag Purification Resin; Roche, USA) for immobilized metal affinity chromatography (IMAC). The resin was incubated with protein at 4 ℃ for 2 hours with gentle mixing and then packed into empty PD10 open columns (Cytiva, USA) for subsequent wash and elution. The wash and elution buffer were supplemented with 3 mM and 250 mM imidazole, respectively. After each purification step, the eluents were checked by SDS-PAGE to identify the target proteins and evaluate their purity. When applicable, the affinity purification tag was removed by incubation with TEV protease at a 1:50 (v/v) ratio overnight at 4 ℃ with simultaneous dialysis to remove residual imidazole. After the final SEC step, individual fractions containing the target proteins were pooled, concentrated by Amicon Ultra centrifugal ultrafiltration filters (Merck Millipore, USA), and checked by a NanoPhotometer N60 (IMPLEN, Germany) to determine their concentrations by their absorbance at 280 nm, aliquoted, snap-frozen and stored at −80 ℃ until further use.

Ub(WT), Ub(K63R), Ub(K43R), and Ub(K11-only) were expressed at 20 ℃ for 16–20 hours after inducing the bacteria cells with 0.6 mM IPTG. The buffer used for cell lysis and subsequent IMAC purification is composed of 30 mM Tris-HCl, pH 7.5, 100 mM NaCl, and 5 mM β-mercaptoethanol (βME). The N-terminal His_6_-tag was removed from the protein by TEV protease followed by IMAC. The remaining impurities and cleaved His-tag were removed in buffer A (30 mM HEPES, pH 7.5, 100 mM NaCl, and 5 mM βME).

The recombinant E1 and the ubiquitin-conjugating enzyme UBCH7 (E2) were expressed at 16 ℃ for 20 hours after inducing the bacteria cells with 0.6 mM IPTG. The harvested cells were disrupted by sonication (Qsonica, USA) in lysis buffer (50 mM Tris-HCl, pH 7.6, 200 mM NaCl, and 5 mM βME).

After IMAC purification, fractions containing the E1 protein were diluted with 50 mM Tris-HCl, pH 7.6, and 2 mM dithiothreitol (DTT) to reduce NaCl concentration to 40 mM and were further separated by anion-exchange chromatography using Q Sepharose Fast Flow resin (Cytiva, USA) packed into an empty PD10 open-column (Cytiva, USA). The E1 protein was eluted with a NaCl step gradient ranging from 0 to 400 mM with a 40 mM increment. The purification was completed by SEC using a Superdex 200 Increase 10/300 column (Cytiva, USA) in buffer B (30 mM HEPES, pH 7.5, 200 mM NaCl, and 0.5 mM TCEP).

The E2 protein construct used for the study contained an N-terminal His_6_-SUMO tag. To remove the SUMO tag, IMAC-purified protein was treated with in-house made His_6_-SENP2 protease at a 1:50 (v/v) ratio. After reducing the imidazole concentration by overnight dialysis at 4 °C, the SUMO tag was removed by Ni-NTA, and the flow-through fractions containing the E2 protein were collected. In the final step, the E2 protein was purified by SEC in buffer B.

The WW3-HECT domain of yeast ubiquitin ligase Rsp5 (E3(WT)) and its mutated version with C-terminal 3 amino acid insertion, namely Rsp5 WW3-HECT_GML (E3), were expressed at 16 °C for 16 hours after inducing the bacteria cells with 0.5 mM IPTG. Cells were lysed in 30 mM Tris-HCl, pH 8.0, 200 mM NaCl, 10% glycerol, 2 mM βME, and 0.5 mM PMSF. The same buffer was also used for washing and elution during subsequent IMAC purification. Both constructs contained an N-terminal His_6_-tag followed by a TEV cleavage site. The target proteins were purified by SEC in buffer C (20 mM HEPES, pH 7.5, 200 mM NaCl, 10% glycerol, and 0.5 mM TCEP).

UCHL5(WT) and UCHL5(C88A) were N-terminally His_6_-tagged. They were over-expressed at 16 °C for 20 hours after induction with 0.4 mM IPTG. Cells were lysed in buffer C (3 mM DTT was used instead of 0.5 mM TCEP). The same buffer was used for the His-tag affinity purification and was supplemented with 3 mM and 200 mM imidazole for wash and elution, respectively. The His_6_-tag was not removed from the expressed protein. They were purified by SEC in buffer C.

RPN13 was N-terminal His-tagged. It was over-expressed in *E. coli* BL21 (DE3) (Yeastern Biotech, Taiwan) strain after induction with 0.4 mM IPTG. Cells were lysed in a buffer containing 20 mM Tris-HCl, pH 7.5, 150 mM NaCl, 5 mM DTT, and 1 mM PMSF. IMAC was carried out in buffer D (20 mM Tris-HCl, pH 8.0, 150 mM NaCl, and 2 mM DTT) supplemented with 5 mM or 250 mM imidazole for wash and elution, respectively. SEC was performed using a HiLoad 16/600 Superdex 75 pg column (Cytiva, USA) in buffer C.

The RPN13-UCHL5 complexes were formed by mixing RPN13 with UCHL5(WT) or UCHL5(C88A) mutant at a molar ratio of 1:1.3. Mixtures were incubated on ice for 2 hours, and excess free protein was removed by SEC in buffer C. The deubiquitinase (DUB) activity of RPN13-UCHL5(WT) was confirmed by comparing the DUB activity of free UCHL5(WT) using a model fluorogenic substrate Ub-AMC (UbpBio, USA)^57^.

Recombinant Sic1^PY^ was overexpressed at 16 °C overnight after inducing the bacteria with 0.1 mM IPTG. The harvested cells were disrupted by sonication (amplitude 10, 10 s/10 s on/off, the total processing time of 20 minutes) in lysis buffer D supplemented with cOmplete™, EDTA-free Protease Inhibitor Cocktail (Roche, USA). Sic1^PY^ was purified by IMAC aided by its C-terminal His-tag. IMAC was performed in 20 mM Tris-HCl, pH 8.0, 200 mM NaCl, 10% glycerol, and 2 mM DTT.

His-tagged Lb^pro^ (29–184 aa, UniProt ID: P05161) was synthesized and subcloned to pRSF-duet expression vector by GenScript (Piscataway, NJ). Lb^pro*^(L102W) was generated by site-directed mutagenesis to cleave ubiquitin^2^. Over-expression of Lb^pro*^ was induced by 0.6 mM IPTG. Lb^pro*^ was purified by IMAC, followed by SEC using a HiLoad 16/600 Superdex 200 pg column (Cytiva, USA). Fractions containing Lb^pro*^ were collected, concentrated, and flash-frozen by liquid nitrogen.

### Site-specific fluorescent labeling of ubiquitin and Sic1^PY^

To facilitate the substrate detection in electrophoretic gels and distinguish deubiquitination of the substrate from Sic1^PY^ proteolysis, we fluorescently labeled Ub with Fluorescein-5-Maleimide (AnaSpec, USA) and Sic1^PY^ with Alexa Fluor 647 C_2_ Maleimide (Thermo Fisher Scientific, USA). A cysteine was engineered at the C-terminus of Sic1^PY^; a cysteine was introduced to the N-terminus of a Ub variant. Both proteins were fluorescently labeled through maleimide-mediated covalent modification. Fluorescein-5-Maleimide and Alexa Fluor 647 C_2_ Maleimide were dissolved according to manufacturer instructions at 0.1 M concentration. Before adding the dye, the protein was reduced by DTT at a final concentration of 30 mM on ice for 30 min. The reducing agent was removed by desalting using a disposable PD 10 desalting column (Cytiva, USA). The reduced protein in 25 mM HEPES, pH 7.0, and 150 mM NaCl was incubated with a five-fold excess of the dye at 4 ℃ overnight. The reaction was stopped by adding a quenching buffer to a final concentration of 50 mM Tris and 20 mM DTT. A second desalting step subsequently removed the unreacted fluorescent dye. The fluorescently labeled proteins were resolved by 4-20% SurePAGE (GeneScript, USA), and the protein concentration was estimated based on the measured dye-specific absorbance.

### Preparation of Sic1^PY^-Ub_n_

*In vitro* ubiquitination assay was performed in a 10 mL reaction mixture containing 50 mM HEPES, pH 7.0, 250 mM NaCl, 10 mM MgCl_2_, 7.5 mM ATP, 0.05 mg/mL BSA, 250 nM UBE1 (E1), 750 nM UbcH7 (E2), 1.2 µM Rsp5-HECT_GML_ (E3), together with 7.15 µM Ub (10% fluorescein-labeled) and 1.25 µM Sic1^PY^ (10% AlexaFluor-647-labelled) at 37 ℃ for 30 min. The reaction mixture was incubated at 4 ℃ for 2 hours with 3 mL of cOmplete Ni-NTA resin (Roche, Germany) equilibrated in buffer E (50 mM Tris-HCl, pH 7.5, 500 mM NaCl, 0.005% NP40). Polyubiquitinated Sic1^PY^ and E1 (both His-tagged) were eluted by buffer E supplemented with 250 mM imidazole. The product was fractionated and E1 removed by SEC using a Superdex 75 increase 10/300 column (Cytiva, USA) with buffer F (30 mM HEPES, pH7.5, 100 mM NaCl, 5% glycerol, 0.5 mM TCEP, and 0.005% NP-40). The fractions containing the substrate with Ub chains of intermediate length (4-10 Ubs) were collected and used for further experiments.

### Native-gel separation of 26S proteasome

To resolve the proteasomal complexes, 3–8 % Tris-acetate gels (Invitrogen, USA) were used for native electrophoresis. The reconstituted samples were mixed with 4× loading buffer (100 mM Tris-HCl, pH 8.0, 20% glycerol, 2 mM ATP, 0.002% bromophenol blue) immediately before loading. 1-2 ug of proteasomal complex was loaded per well. The separation was carried out at 4℃ and 150 V for 4 hours, using a TBE running buffer supplemented with 0.5 mM DTT, 0.5 mM ATP, and 2 mM MgCl_2_ in a XCell SureLock Mini-Cell (Invitrogen, USA).

### In-gel 26S proteasome activity

Native gels containing 26S proteasomes were incubated with reaction buffer (50 mM Tris-HCl, pH 7.5, 10 mM MgCl_2_, 1 mM ATP, 1 mM DTT, 50 μM Suc-LLVY-AMC) for 30 min at 37 ℃.

The fluorescence signal of the product generated by 26S proteasomes around the complex position in the gel was visualized and quantified by ChemiDoc XRS+ Imaging Systems with supplied Image Lab Software (BioRad, USA).

### Immunodetection of native 26S proteasome components

For immunoblotting, the native gels containing the 26S proteasomes were transferred to Amersham Hybond LFP, 0.2 PVDF membranes (Cytiva, USA) using Bio-Rad mini-protean 3 transfer system and buffer containing 25 mM Tris base, 192 mM glycine, 0.1% SDS (sodium dodecyl sulfate) for 4 hours at 70 V (current limit: 350 mA) in cold room. The membranes were incubated with blocking buffer containing 5% non-fat dry milk in TBST (Tris-buffered saline with 0.1% Tween-20) at room temperature for 1 hour and then incubated with primary antibodies diluted in blocking buffer at 4 ℃ overnight. Next, the membranes were washed five times for 5 min each with TBST and then incubated with horseradish peroxidase (HRP)-conjugated secondary antibodies ×10,000-fold diluted in blocking buffer at room temperature for 1 hour. After five times 5 min wash with TBST, the membrane was covered with Clarity Western ECL Blotting Substrate (BioRad, USA) and incubated for 5 min. Finally, the developed HRP signal was recorded by iBright Imaging System (Invitrogen, USA).

For re-probing with different antibodies, the membranes were stripped by incubating the membrane five times for 10 min each in a solution containing 1.5% glycine, pH 2.2, 0.1% SDS, and 1% Tween20. The membrane was subsequently washed twice for 10 min each with PBS and twice for 10 min each with TBST. The membrane was then again blocked, and proteins were detected in the same way as described above.

### Western blot analysis of ubiquitinated substrates

The ubiquitinated substrates were resolved by a 4-20% SurePAGE (GeneScript, USA) using Tris-MOPS running buffer supplied with the gels. Before loading on the gel, samples were mixed with 4× loading buffer (200 mM Tris–HCl (pH 6.8), 8% SDS, 0.2% bromophenol blue, 40% glycerol, 20% βME) and heated at 70 ℃ for 10 min. The electrophoretic separation was carried out at room temperature in an XCell SureLock Mini-Cell (Invitrogen, USA) at 200 V for 30 min. Gels containing the resolved samples were transferred to nitrocellulose membranes using a Bio-Rad mini-protean 3 transfer system and buffer containing 25 mM Tris base, 190 mM glycine, 20% Methanol for 1 hour at 100 V (current limit: 350 mA) at room temperature. Alternatively, the proteins were transferred to a PVDF membrane using iBlot Dry Blotting System and dedicated transfer stacks (Invitrogen, USA). The immunodetection procedure was performed the same way described above for analysis of the native 26S proteasome complexes.

### Negative stain electron microscopy (NSEM)

Four microliters of 30× diluted human 26S proteasome (1 mg/mL, UBPBio, USA) alone or mixed with SEC-purified ubiquitinated substrate were placed on carbon-coated grids that were glow-discharged at 25 mA for 30 s. After staining with 2% uranyl formate, the grids were blotted and dried in the air for 1 day. Images were collected using a FEI Tecnai G2-F20 electron microscope at 200 keV (FEI, USA). A magnification of 29,000× was used, corresponding to a pixel size of 2.9 Å. The micrographs were converted to mrc format using EMAN2 and imported to Relion 3.0^58^ for particle picking. Picked particle stacks were exported to cryoSPARC v2.14^59–61^, where further data processing steps were conducted, including 2D classification, *ab-initio* 3D reconstruction, 3D classification, and final 3D refinement. The resulting 3D structures were visualized by using UCSF-ChimeraX^62,63^.

### Mass spectrometry analysis of Ub chain compositions

Sic1PY-Ubn was treated with Lbpro* to determine the polyubiquitin chain architecture. A batch of the ubiquitinated substrate was prepared following the above ubiquitination protocol without fluorescently labeled Sic1^PY^ and Ub. The purified Sic1^PY^-Ub_n_ was treated with 10 µM Lb^pro*^ in 25 mM Tris-HCl, pH 7.5, 100 mM NaCl, and 10 uM EDTA and incubated at 37 ℃ for 20 hours. To determine Ub linkage types, the Lb^pro*^-treated sample was separated by 4-20 % SurePAGE (GeneScript, USA) using Tris-MOPS running buffer supplied with the gels and stained with One-Step Blue^®^ (Biotium, USA). The protein bands were excised from the gel, diced, and destained with 25% (v/v) acetonitrile (ACN) and 5 % Trifluoroacetic acid (TFA). The diced gel was reduced by 50 mM Dithioerythritol (DTE) in 25 mM ammonium bicarbonate (ABC) at 37 ℃ for 1 hour, and subsequently alkylated by 100 mM iodoacetamide (IAM) at room temperature in the dark for 1 hour. The diced gel was washed with 25% (v/v) ACN in 25 mM ABC, pH 8.5 for 5 min once, dehydrated with 25% ACN in 25 mM ABC for 5 min, and dried under vacuum. The gel pieces were rehydrated with a 1:20 protease to protein ratio with a Mass Spectrometry Grade Lys-C (Fujifilm-Wako, USA) in 25 mM ABC digested at 37 ℃ for 3 hours. Following Lys-C digestion, the same amount of sequencing grade Trypsin (Promega, USA) in 25 mM ABC was added for digestion at 37 ℃ for 16 hours. Tryptic peptides were extracted twice with 50% (v/v) ACN containing 5% (v/v) TFA for 3 min each time with moderate sonication. The extracted solutions were pooled and dried by speed vacuum. The peptide mixture was aliquoted and desalted by a C18-ZipTip (Merck Millipore, USA) and eluted with 50% ACN in 0.1% (v/v) formic acid (FA). 200 fmol of ubiquitin-absolute quantification (Ub-AQUA) peptide, which is a gift from Dr. Yasushi Saeki in Tokyo Metropolitan Institute of Medical Science, Japan, was added into the extracted peptide as ideal internal standards and subsequently concentrated peptide mixture to a final volume of less than 2 µL using by speed vacuum followed by H_2_O_2_ treatment by diluting the peptide mixture to 20 microliters of 0.1 % TFA and 0.05 % H_2_O_2_ at 4 ℃ overnight to oxidize methionine. Finally, the ubiquitin-absolute quantification/ parallel reaction monitoring (Ub-AQUA/ PRM) assay was used to estimate the abundance of each Ub-chain type linkage of signature peptides with di-glycine (GG) modification at a specific lysine against the total number of the modified peptides detected in the sample.

LC-MS-based Ub-AQUA/ PRM analysis was performed on a Thermo UltiMate 3000 RSLCnano system connected to a Thermo Orbitrap Fusion mass spectrometer (Thermo Fisher Scientific, Germany) equipped with a nanospray interface (New Objective, USA). Peptide mixtures were loaded onto a 75 μm ID, 25 cm length C_18_ BEH column (Waters, USA) packed with 1.7 μm particles with a pore of 130 Å and were separated using a segmented gradient in 30 min from 5% to 35% solvent B (0.1% (v/v) FA in ACN) at a flow rate of 300 nL/min. Solvent A was 0.1% (v/v) FA in water. The mass spectrometer was operated in the PRM mode. Briefly, the MS^1^ scans of peptide precursors from 350 to 1600 m/z were performed at 120 K resolution with a 2 × 10^5^ ion count target. The PRM scans were performed by isolation window at 1 Da with the quadrupole, HCD fragmentation with normalized collision energy of 30, and analysis at 30 K resolution in the orbitrap. The PRM scans ion count target was 1 × 10^6,^ and the max injection time was 54 ms. The PRM scans were triggered by time-scheduled targeting precursor ions selected for target peptides (without heavy isotope labeling, Supplementary Table 5) in ±2 min elution windows.

The PRM data were processed using Skyline 22.2.0.255 (MacCoss Lab Software, USA) to generate XIC and perform peak integration. All peaks were manually inspected to confirm correct detection and peak boundaries. Peak integration and calculation of the ratios between light endogenous and the heavy-labeled peptide (L/H) were done in Skyline, and result reports were exported from the software. Light-to-heavy peptide peak area ratios were used for the target peptide quantitation analysis.

For determining the branching of the Ub chains, an aliquot of the Lb^pro*^-treated substrate sample corresponding to two micrograms of the pure protein was first diluted with 50% (v/v) Methanol followed by 50% (v/v) ACN and 1% (v/v) FA. An aliquot corresponding to one pmol of the pure protein was subsequently injected via the LockSpray Exact Mass Ionization Source (Waters, USA) with a syringe pump (Harvard Apparatus, USA) and held a flow rate of 3 uL/min throughout the analysis to the mass spectrometer. The mass of intact proteins was determined using Waters Synapt G2 HDMS mass spectrometer (Waters, USA). The acquired spectra were deconvoluted to zero-charge spectrum using MaxEnt1 algorithm of the MassLynx 4.1 software (Waters, USA).

### Cryo-EM sample preparation

To prepare 26S-UCHL5(C88A)-Sic1^PY^-Ub_n_(K63R) complexes, 40 μL of 1 mg/mL purified human 26S proteasome (UBPBio, USA) was first incubated at 37 ℃ for 10 min and then supplemented with preformed RPN13:UCHL5(C88A) to a final concentration of 1 μM and incubated for another 10 min at room temperature. To prevent substrate *en bloc* deubiquitination by RPN11, the sample was treated with 3 mM o-phenanthroline (oPA) and incubated for an extra 5-10 min. Next, 40 μL of 2-3 µM ubiquitinated Sic1^PY^ substrate, supplemented with 2 mM ATP and 5 mM MgCl_2_, was added to the mixture and incubated for another 5 min at room temperature. The sample was then diluted with 350 μL of cold buffer composed of 20 mM HEPES, pH 7.6, 40 mM NaCl, 5 mM MgCl_2_, 2 mM βME, 0.5 mM oPA, and concentrated to 40 μL using 100 kDa MWCO Amicon Ultra-0.5 mL Centrifugal Filter (MerckMillipore, USA) to reduce glycerol content. The sample was incubated at room temperature immediately before vitrification. Three microliters of the reconstituted sample were applied onto 300-mesh Quantifoil R1.2/1.3 holey carbon grids, which were glow-charged at 20 mA for 30 sec before use. After incubation for 3 s, the grids were blotted for 2.5 s at 4 ℃ and 100% humidity and vitrified using a Vitrobot Mark IV. (Thermo Fisher Scientific, USA).

### Cryo-EM data collection

Cryo-EM data acquisition was performed on a 300 keV Titan Krios transmission electron microscope (Thermo Fisher Scientific, USA) equipped with a Gatan K3 direct detector (Gatan, USA) in a super-resolution mode using the EPU software (Thermo Fisher Scientific, USA). Movies were collected with a defocus range of −1.2 to −1.8 μm at a magnification of 64,000×, corresponding to a super-resolution pixel size of 0.7 Å. A total of 48–50 e^-^/Å^2^ was distributed over 50 frames with an exposure time of 1.8 s. The datasets were energy-filtered with a slit width of 15–20 eV, and the dose rates were adjusted to 8–10 e^-^/pixel/s.

### Cryo-EM data processing and model building

A total of 15,242 micrograph movies were recorded, then aligned and two-fold binned (resulting in a pixel size of 1.4 Å) using MotionCor2 (UCSF)^64^ and CTF corrected by CTFFIND4.1^65^. The particle images of the single-(RP-CP) and double-capped (RP-CP_2_) proteasome particles were manually picked and used as a training set for automated particle picking by Topaz convolutional neural network, which resulted in an initial dataset of 2,695,911 particles^66^. Further data processing, including 2D and 3D classifications, was performed in RELION-3.1^67^, while the final nonuniform refinement and variability analysis was performed in CryoSparc 3.1 (Structura Biotech, Canada)^59–61^. The initial dataset of picked particles was extracted, and four-fold binned (resulting in a pixel size of 5.6 Å, box size of 150 pixels) was cleaned by 2D and global 3D classification, resulting in a dataset containing 575,778 RP-CP and 527,610 RP-CP_2_ proteasome particles. The particles representing RP-CP_2_ due to their pseudo symmetry were refined, imposing the C2 symmetry restrain, then symmetry-expanded (doubling the number of the RP-CP_2_ particles), and again refined with C1 symmetry with mask focused on only one RP of the complex. The symmetry expansion procedure was used to allow the merging of the RP-CP_2_ particles with the RP-CP complexes while simultaneously reducing the heterogeneity of the dataset. After combining, all the particles were re-centered on one RP (RP of interest, in the case of the symmetry-expanded RP-CP2 the refined RP) and re-extracted with two-fold binning (resulting pixel size of 2.8 Å, box size of 300 pixels). After 3D classification focused on the RP of interest, the particles’ signal was partially subtracted, leaving only the signal of the RP of interest and the first ring of α subunits composing the CP subcomplex. The classification yielded two conformations of the RP, one with 532,818 particles resembling E_D_ active state of the human 26S complex and the second with 736,915 RPs resembling the resting E_A_ state of the human 26S proteasome. After global refinement, particles in each subset were re-extracted at full resolution (resulting in a pixel size of 1.4 Å, box clipped to 400 pixels). Further classifications of the E_A_-like particles focused on the ubiquitin-binding region filtered out 153,646 particles without any obvious proteasome-bound ubiquitin densities. The subset produced 3.4 Å resolution map of “empty” proteasomes after nonuniform refinement in Cryosparc3.1. Heterogeneity among the remaining 202,823 E_A_-like particles was resolved by 3D Variability Analysis within Cryosparc3.1 (Structura Biotech, Canada)^59–61^, which revealed the existence to E_A_-like proteasomes with three (E_A_:Ub_3_) or E_B_-like proteasomes with four (E_B_:Ub_4_) ubiquitin bound at the RP, which accounted for 54,794 and 52,216 particles, respectively. The particles representing both states were then refined by nonuniform refinement in Cryosparc3.1, yielding a map of the E_A_:Ub_3_ and E_B_:Ub_4_ with an overall resolution of 3.6 Å and 3.8 Å, respectively.

The subset of E_D_ particles was also classified with a mask focused on the ubiquitin-binding region. Traces of proteasome-bound Ub were found in all of the obtained classes; however, for further processing, only 212,581 particles with the best and unambiguously defined Ub densities were selected. Based on subsequent classification aimed at resolving heterogeneity in the AAA+ ATPase, we extracted two subsets of E_D_:Ub_4_ particles with slightly different conformations of the ATPase subcomplex accounting for 84,476 and 49,828 particles, which, after nonuniform refinement in Cryosparc3.1 produced maps with an overall resolution of 3.8 Å and 4 Å, respectively. Both of the reconstructed E_D_:Ub_4_ maps did not differ significantly in the region of interest, i.e., the Ub and Ub-binding region and the 3.8 Å showed better details of the AAA+ ATPase we used this map for further building of the atomic model of the ED:Ub4 state. All the reconstructed cryo-EM maps of the Ub-bound complexes were further focus-refined with masks around RPN3:RPN7 region, Ub and neighboring interfaces, and AAA+ ATPase. The obtained focused maps were merged in Phenix using Combine Focused Maps procedure^68^. The resulting combined cryo-EM maps were used for final atomic model refinement. The atomic models of the empty complex, E_A_:Ub_3_, E_B_:Ub_4_, and E_D_:Ub_4_ states of the complex were built using Phenix and COOT^68,69^ based on PDB 6MSB (E_A_ state) and 6MSK (E_D_ state) structures as references^37^. Atomic details and interactions of the proteasome-bound Ub K11/K48 chain and the neighboring subunits were refined using ISOLDE^70^ within UCSF ChimeraX^62,63^. The latter was also used to visualize the cryo-EM maps and atomic models. All cryo-EM data collection and refinement statistics are reported in Supplementary Table 4.

### Substrate degradation assay

The rate and efficiency of proteasome-mediated degradation of Sic1^PY^ ubiquitinated with branched Ub chains (generated using Ub(K63R) mutant), or with homotypic K11-, or K48-linked chains (generated using the Ub(K11-only) or Ub(K48-only) variants, respectively) was estimated based on quantification of the degradation products separated in SDS-PAGE. The degradation reaction contained 60 nM human 26S proteasome in 30 mM HEPES, pH 7.4, 50 mM NaCl, 10% glycerol, 0.5 mM TCEP, 0.5 mg/mL BSA, ATP regeneration system (50 mM ATP, 100 mM MgCl_2_, 3.2 mg/mL creatine kinase, 160 mM creatine phosphate), and 10 nM of Sic1^PY^-Ub_n_. The concentrations of the substrates labeled with branched-, K11-, and K48-linked Ub chains used for the degradation assay were normalized based on the AlexaFluor-647 signal of each of the substrate preparations. To check the effect of UCHL5 on the proteasome-mediated degradation of the substrates, the reactions were additionally supplemented with preformed SEC-purified complexes of RPN13-UCHL5(WT), catalytically inactive RPN13-UCHL5(C88A), or RPN13 alone as a control. The reactions were carried at 37°C for 30 min, after which they were stopped by adding SDS-PAGE loading dye into the sample and 10 min heating at 37 ℃. Degradation products and remaining undegraded substrates were resolved by a 4-20% SurePAGE (GeneScript, USA) using Tris-MOPS running buffer supplied with the gels. The amount of degraded substrate was estimated by quantifying the AlexaFluor-647 signal of the small molecular weight degradation products appearing in the very bottom of the gel in the samples treated with the proteasome. The degradation efficiency of each of the tested Ub chains (expressed as a faction of degraded substrate labeled with the particular Ub chain) was calculated by comparing the signal of the degradation products with the total signal of the degradable substrate, i.e., the sum of remaining substrate labeled with Ub chains of at least four Ub (corresponding to signals of molecular weight of ≥50 kDa) and the degraded substrate (which equals to the signal of degradation product). The statistical significance of the observed differences was calculated using two-way ANOVA performed using GraphPad Prism 8 (GraphPad Software, USA).

### Sample Preparation and Cross-Linking

The functional 26S proteasome complex samples for the XL-MS experiment were prepared the same way as for the cryo-EM experiment, except that one of the samples was prepared without the substrate. The formed complex samples, with and without the substrate, each containing 50 μg of the human 26S proteasome (UBPBio, USA) in a total volume of 100 μL were then individually cross-linked using disuccinimidyl sulfoxide (DSSO) at a final concentration of 80 μM for 1 hour. The reactions were stopped by adding ABC to a final concentration of 5 mM.

Cross-linked samples were then buffer-exchanged to 50 mM triethylammonium bicarbonate buffer using 100 kDa MWCO Amicon Ultra-0.5 mL Centrifugal Filter (Merck Millipore, USA). Samples were reduced and alkylated by 10.7 mM TCEP and 21.4 mM chloroacetamide, then incubated at 95 ℃ for 10 min. Proteins were proteolyzed with Lys-C (1:50 w/w) and trypsin (1:50 w/w) for 4 hours at 37 ℃. The peptide digests were desalted using Sep-Pak C8 cartridges (Waters, USA), dried under vacuum, and reconstituted in 0.05% TFA and 1% (v/v) ACN for subsequent LC/MS measurements.

Samples were separated by reverse phase LC using a Thermo Scientific™ UltiMate™ 3000 UHPLC system connected to a Thermo Scientific™ EASY-Spray™ column, 50 cm × 75 µm over a 130 min with 4-25% gradient (A: water, 0.1% FA; B: ACN, 0.1% FA) at a flow rate of 300 nL/min. The cross-linked samples were analyzed on a Thermo Scientific™ Orbitrap Eclipse™ Tribrid™ mass spectrometer with FAIMS Pro™ device (Thermo Fisher Scientific, USA) and Instrument Control Software version 3.4 using a stepped collision energy (SCE) MS^2^ HCD of 21-26-31% or MS^2^CID-MS^3^CID acquisition strategies. MS^2^ sequencing events were triggered for precursors of charge states +3 to +8. Mass-difference-dependent CID-MS^3^ acquisitions were triggered if a unique mass difference (Δ = 31.9721 Da) was observed in the CID-MS^2^ spectrum. MS^1^ and MS^2^ scans were acquired in the Orbitrap with a mass resolution of 60,000 and 30,000 correspondently, whereas MS^3^ scans were acquired in the ion trap. Precursor isolation windows were set to 1.6 m/z for MS^2^ scans and 2.5 m/z for MS^3^ scans.

### Cross-linking identification and quantification

The raw data were analyzed using Proteome Discoverer 2.5 software (Thermo Fisher Scientific, USA) with XlinkX node 2.5 for cross-linked peptides and a SEQUEST HT search engine for unmodified peptides and monolinks. The following parameters were applied for Proteome Discoverer: MS^1^ ion mass tolerance, 10 ppm; MS^2^ ion mass tolerance, 20 ppm; MS^3^ ion mass tolerance, 0.6 Da; maximal number of missed cleavages, 2; minimum peptide length, 6.

Carbamidomethylation (+57.021 Da) was used as a static modification for cysteine. Different cross-linked mass modifications were used as variable modifications for lysine in addition to methionine oxidation (+15.996 Da). Data were searched against a database containing Uniprot/SwissProt entries of human SwissProt database (retrieved Oct. 2020) based on protein identifications from the proteomics experiment of 26S proteasome complex. The false discovery rate (FDR) was set to 1% at the CSM level and 5% at the cross-linkers level. Additionally, cross-links were filtered with an identification score of ≥40 and Δ-score of ≥4. Results visualization was performed using xiNET2 v18.3.23.

## Data and materials availability

The atomic coordinates of the apo E_A_ state generated in this study have been deposited in the Protein Data Bank (PDB) under the accession code 8JRI (E_A_ state). The atomic coordinates of the 26S proteasome complexed with substrates in different states generated in this study have been deposited in the Protein Data Bank (PDB) under the accession codes 8JRT (E_A_:Ub_3_ state), 8JTI (E_B_:4Ub_4_ state) and 8K0G (E_D_:Ub_4_ state). The cryo-EM map of the apo E_A_ state has been deposited in the Electron Microscopy Data Bank (EMDB) under the accession codes EMD-36598 (consensus). The cryo-EM maps of the E_A_:Ub_3_ state have been deposited in the EMDB under the accession codes EMD-36605 (composite), EMD-37269 (consensus), EMD-37273 (local refinement focused on Ub binding region), EMD-37276 (local refinement focused on Rpn3/Rpn7 region), and EMD-37277 (local refinement focused on AAA+ ATPase subcomplex). The cryo-EM maps of the E_B_:Ub_4_ state have been deposited in the EMDB under the accession codes EMD-36645 (composite), EMD-37317 (consensus), EMD-37319 (local refinement focused on Ub binding region), EMD-37327 (local refinement focused on Rpn3/Rpn7 region), and EMD-37328 (local refinement focused on AAA+ ATPase subcomplex). The cryo-EM maps of the E_D_:Ub_4_ state have been deposited in the EMDB under the accession codes EMD-36764 (composite), EMD-37334 (consensus), EMD-37335 (local refinement focused on Ub binding region), EMD-37341 (local refinement focused on the RP lid), and EMD-37344 (local refinement focused on AAA+ ATPase subcomplex). The RAW files of XL-MS analysis have been deposited to MassIVE under the accession codes MSV000092664 (Cross-link MS analysis of the human 26S proteasome in complex with RPN13:UCHL5 without Sic1^PY^-Ub_n_), MSV000092665 (Cross-link MS analysis of human 26S proteasome in complex with RPN13:UCHL5 with Sic1^PY^-Ub_n_), MSV000092883 (Ub-AQUA/ PRM) and MSV000092884 (Intact MS).

## Acknowledgements

We thank Dr. Yasushi Saeki at the Tokyo Metropolitan Institute of Medical Science, Japan, for sharing the Ub-AQUA peptides. We thank Academia Sinica Common Mass Spectrometry Facilities (AS-CFII-111-209) and Academia Sinica Cryo-EM Center (AS-CFII-111-210) for data collection; both are funded by the Academia Sinica Core Facility and Innovative Instrument Project. Taiwan Protein Project (AS-KPQ-109-TPP2) is acknowledged for supporting the cryo-EM facility. We thank the protein core facility, biophysics core facility and imaging core facility of the Institute of Biological Chemistry, Academia Sinica, for supporting the sample preparation and analyses.

## Author contributions

Conceptualization: PD, STDH

Methodology: PD, SYL, RV, YCC, KPW, STDH

Investigation: PD, SNC, TC, YSW, JYCH, RV, STDH

Visualization: PD, TC, YSW, STDH

Funding acquisition: STDH

Project administration: PD, STDH

Supervision: PD, STDH

Writing – original draft: PD, STDH

Writing – review & editing: PD, TC, YSW, STDH

## Supplementary Information

Supplementary Text

Supplementary Figures. 1 to 13

Supplementary Tables1 to 5

Supplementary Movie 1

