## Supporting information for "Structural basis of K11/K48-branched ubiquitin chain recognition by the human 26S proteasome"

**The PDF file includes:**

Supplementary Text
Supplementary Figures 1 to 13
Supplementary Tables 1 to 5

**Other Supplementary Information for this manuscript include the following:**

Supplementary Movies 1

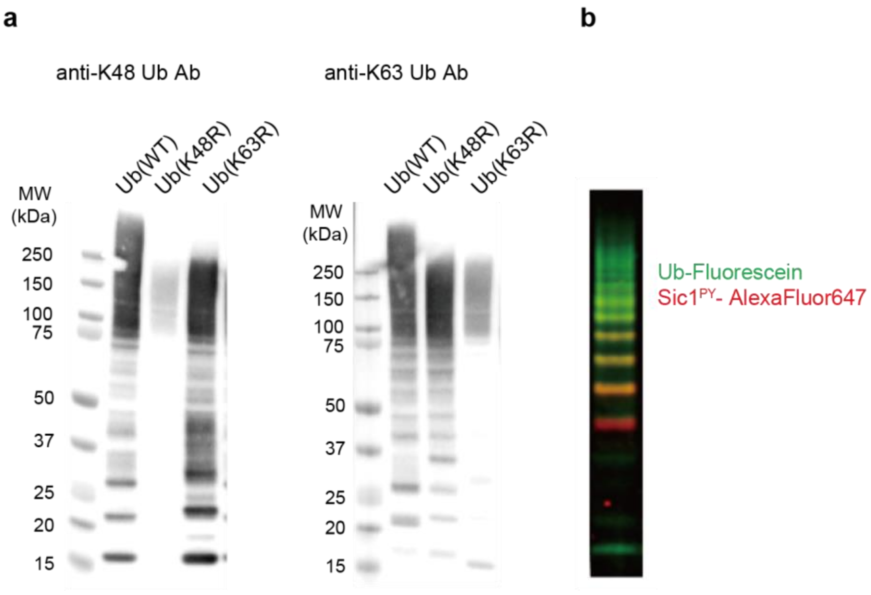

**Supplementary Fig. 1. Characteristics of the ubiquitinated Sic1$^{PY}$ substrate. a**, Use of Ub(K63R) prevented the formation of unwanted K63-linked Ub conjugates. The opposite was true when Ub(K48R) was used in the ubiquitination assay. Anti-K48-linked and anti-K63-linked Ub antibodies were used to probe the linkage types, which are indicated above. **b**, Two-color fluorescence probing of the ubiquitination product using fluorescein-labeled Ub (green) and AlexaFluor647-labeled Sic$^{PY}$ (red). The SDS-PAGE was imaged in two separate channels at the same time to image Ub-Fluorescein and Sic$^{PY}$-AlexaFluor647-labeled.

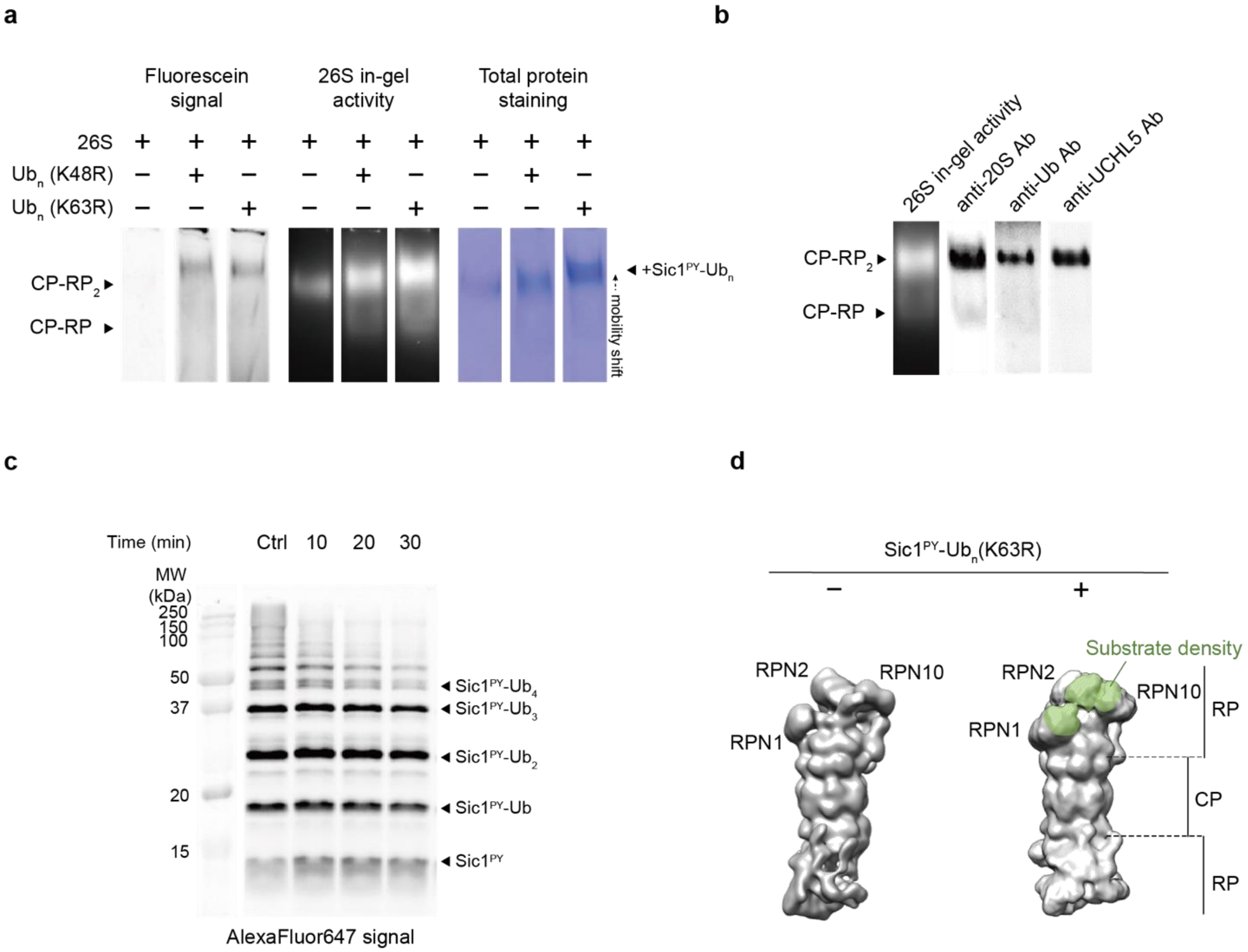

**Supplementary Fig. 2. Validation of the functional 26S complex formation. a,** Native gel electrophoresis detected by fluorescein signals of Ub, in-gel proteolytic activity, and Coomassie Blue staining showed distinct changes in the mobility of the 26S proteasome upon the addition of $Sic1^{PY}$-$Ub_n$(K63R) or $Sic1^{PY}$-$Ub_n$(K48R). **b**, Immunodetection of the native gel-separated samples confirmed the formation of the tertiary complex of 26S-$Sic1^{PY}$-$Ub_n$(K63R)-UCHL5(C88A). **c**, Kinetics of $Sic1^{PY}$-$Ub_n$(K63R) degradation by 26S proteasome monitored by the AlexaFluor647 signal of $Sic1^{PY}$. **d**, EM maps of apo and substrate-bound 26S proteasome derived from NSEM. The additional EM map density in the substrate-bound 26S was highlighted in green.

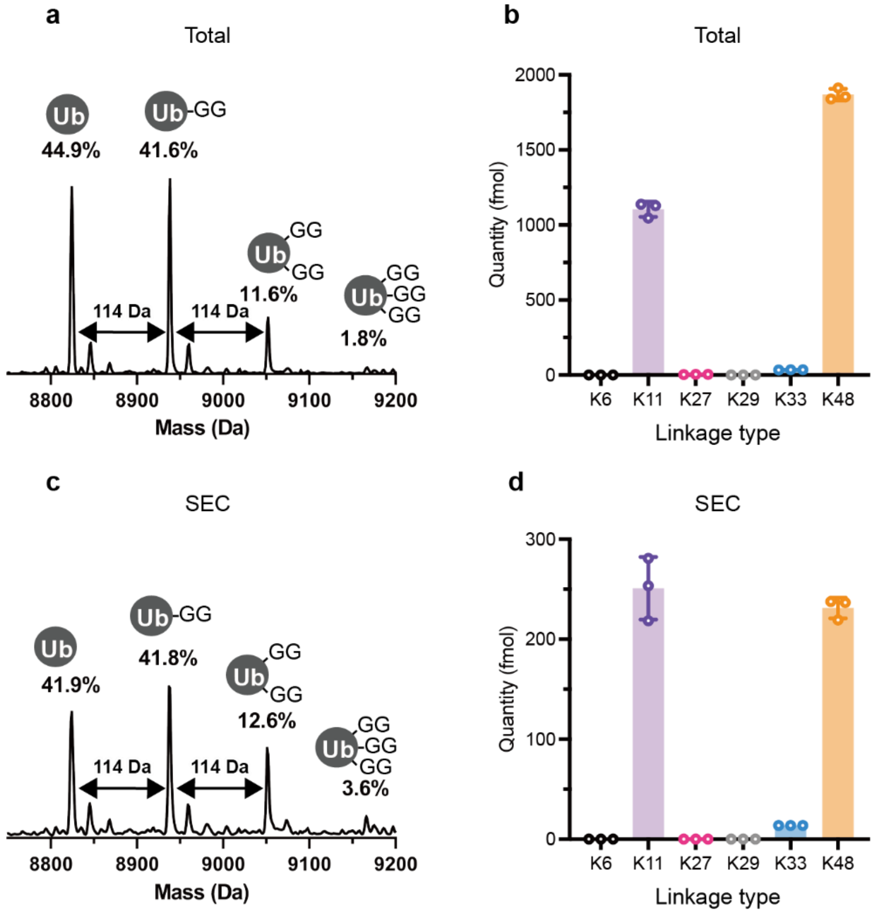

**Supplementary Fig. 3. MS-based quantitative analyses of Ub branching and linkage type distributions.** The amount of branching was quantified by intact MS analysis using $Lb^{pro*}$-digested $Sic1^{PY}$-$Ub_n$. Comparison of the intact mass spectra of total (**a**) and SEC-enriched (**c**) $Sic1^{PY}$-$Ub_n$ showed an increased amount doubly and triply branched Ub after the SEC enrichment. The quantities of the individual peaks were estimated by integrating the areas under the curve. Ub-AQUA analyses of total (**b**) and SEC-enriched (**d**) $Sic1^{PY}$-$Ub_n$. A significant increase of K11-linked Ub was found after SEC enrichment. The experiments were carried in triplicate with the original data points shown in open circles and the error bars indicating the standard deviations.

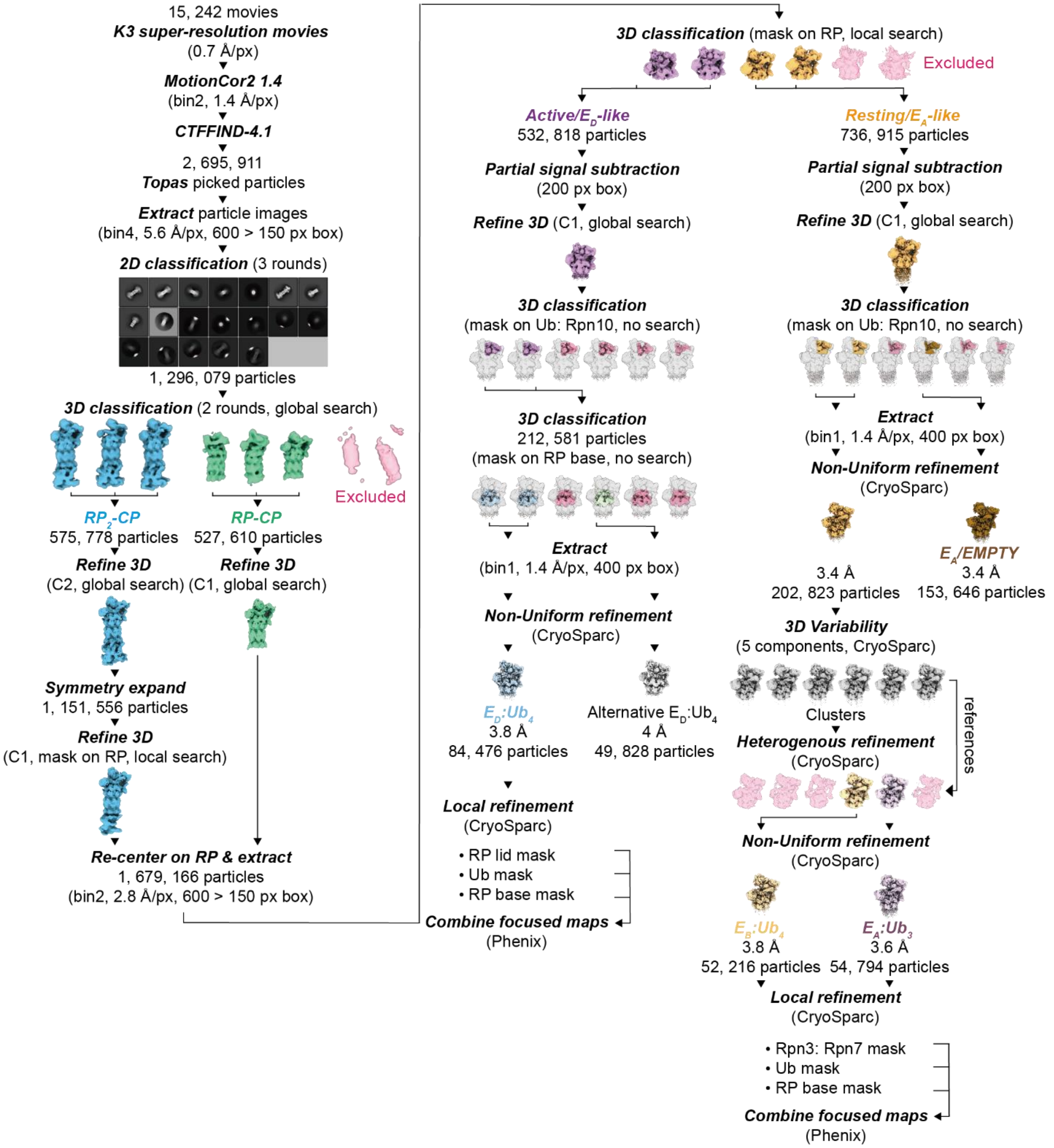

**Supplementary Fig. 4. Cryo-EM data processing pipeline.**

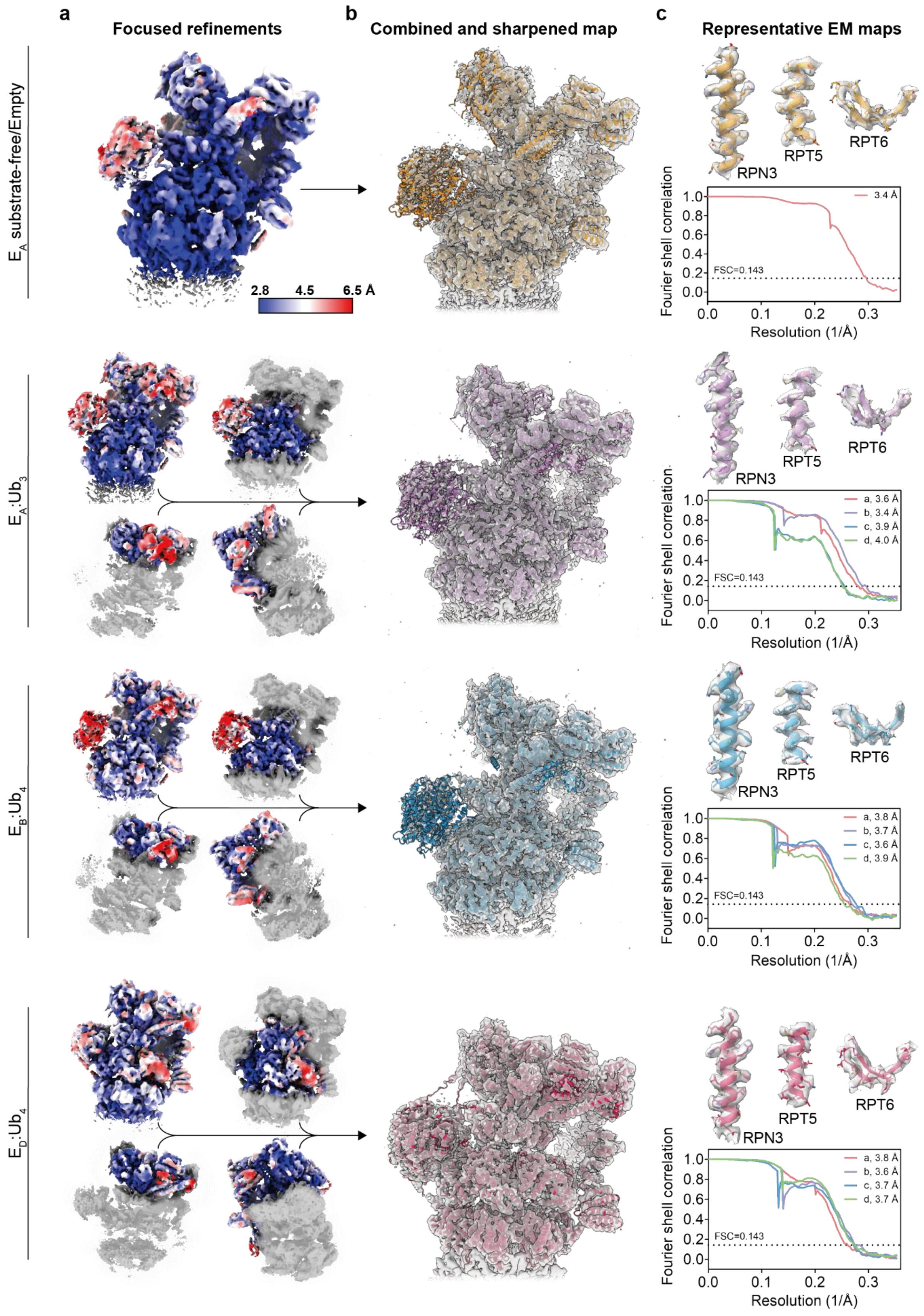
a
Focused refinements
b
Combined and sharpened map
c
Representative EM maps
E_A substrate-free/Empty
2.8
4.5
6.5 Å
RPN3
RPT5
RPT6
Fourier shell correlation
Resolution (1/Å)
FSC=0.143
3.4 Å
E_A:Ub_3
a, 3.6 Å
b, 3.4 Å
c, 3.9 Å
d, 4.0 Å
E_B:Ub_4
a, 3.8 Å
b, 3.7 Å
c, 3.6 Å
d, 3.9 Å
E_D:Ub_4
a, 3.8 Å
b, 3.6 Å
c, 3.7 Å
d, 3.7 Å

71

**Supplementary Fig. 5. Quality assessment of cryo-EM maps. a**, Local resolution estimation of the individual cryo-EM maps of different functional states as indicated on the left. **b**, The composite cryo-EM maps were fused and sharpened (except the $E_A$ substrate-free state) using Phenix Combine Focused Maps and Map Sharpening tools. **c**, Representative cryo-EM maps superimposed with the structural models with the Furrier Shell Correlation curves shown below.

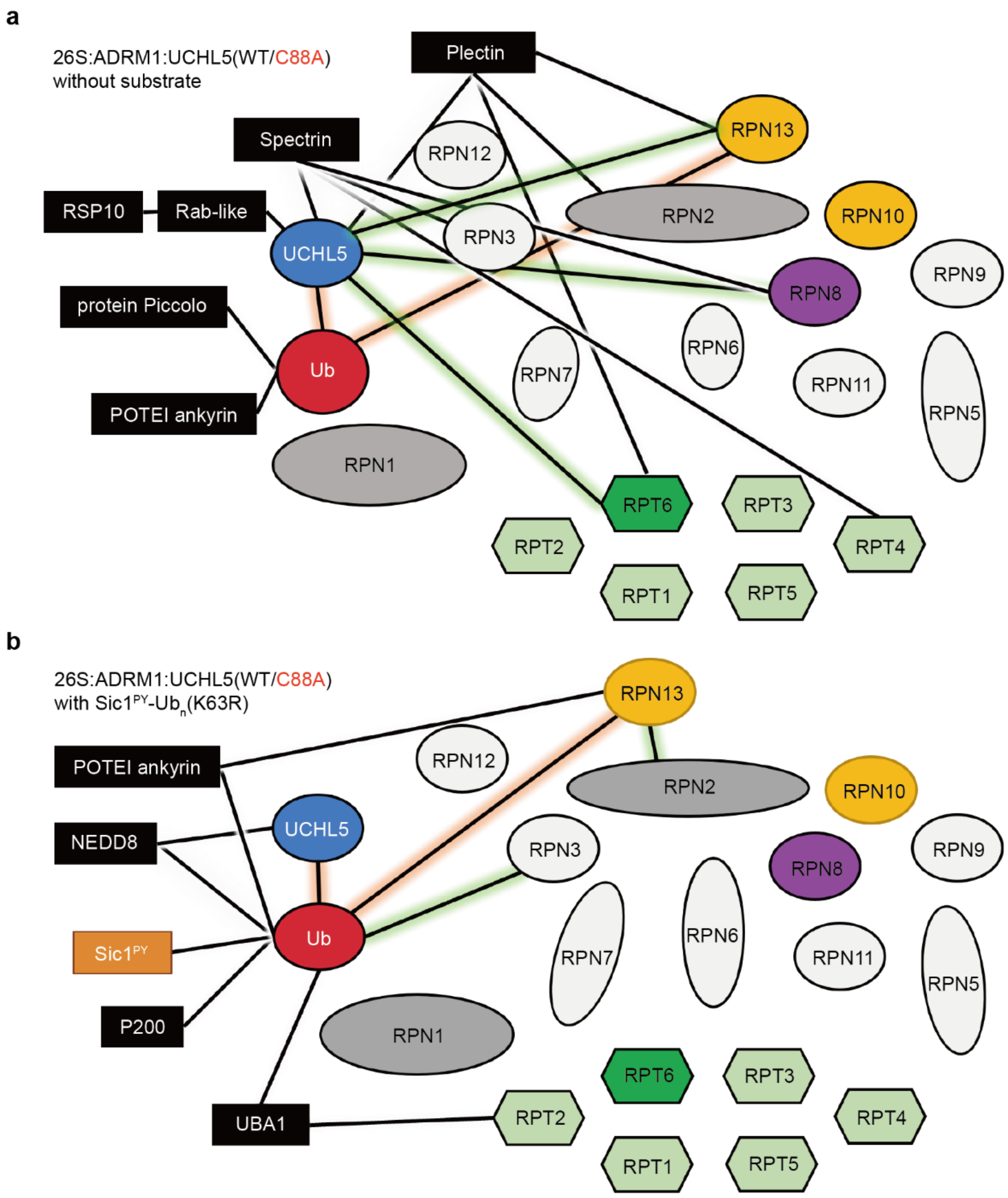

**Supplementary Fig. 6. Interaction network associated with UCHL5 and the 26S proteasome deduced from XL-MS analysis.** The pair-wise protein-protein interactions observed by XL-MS without (**a**) and with the addition of Sic1$^{PY}$-Ub$_n$ (**b**) are shown in black line to link a pair of proteins whose identities are indicated. In the absence of Sic1$^{PY}$-Ub$_n$, UCHL5 was found to interact with several proteasomal subunits, including RPT6 and RPN8. The addition of Sic1$^{PY}$-Ub$_n$ resulted in the loss of these interactions, leading to the only interaction with Ub, suggesting that the Ub chain binding displaced UCHL5 from interacting with other proteasomal subunits. The salmon- and green-shaded lines indicate the cross-links involving UCHL5:RPN13 that were present in both samples or only in one of them. Despite lacking a defined EM map to locate UCHL5 on the proteasome, XL-MS analysis confirmed the presence of UCHL5 on the 19S RP. At the same time, many endogenous proteins were also implicated in the interaction network, but their relative quantities were too low to be resolved by cryo-EM data analysis.

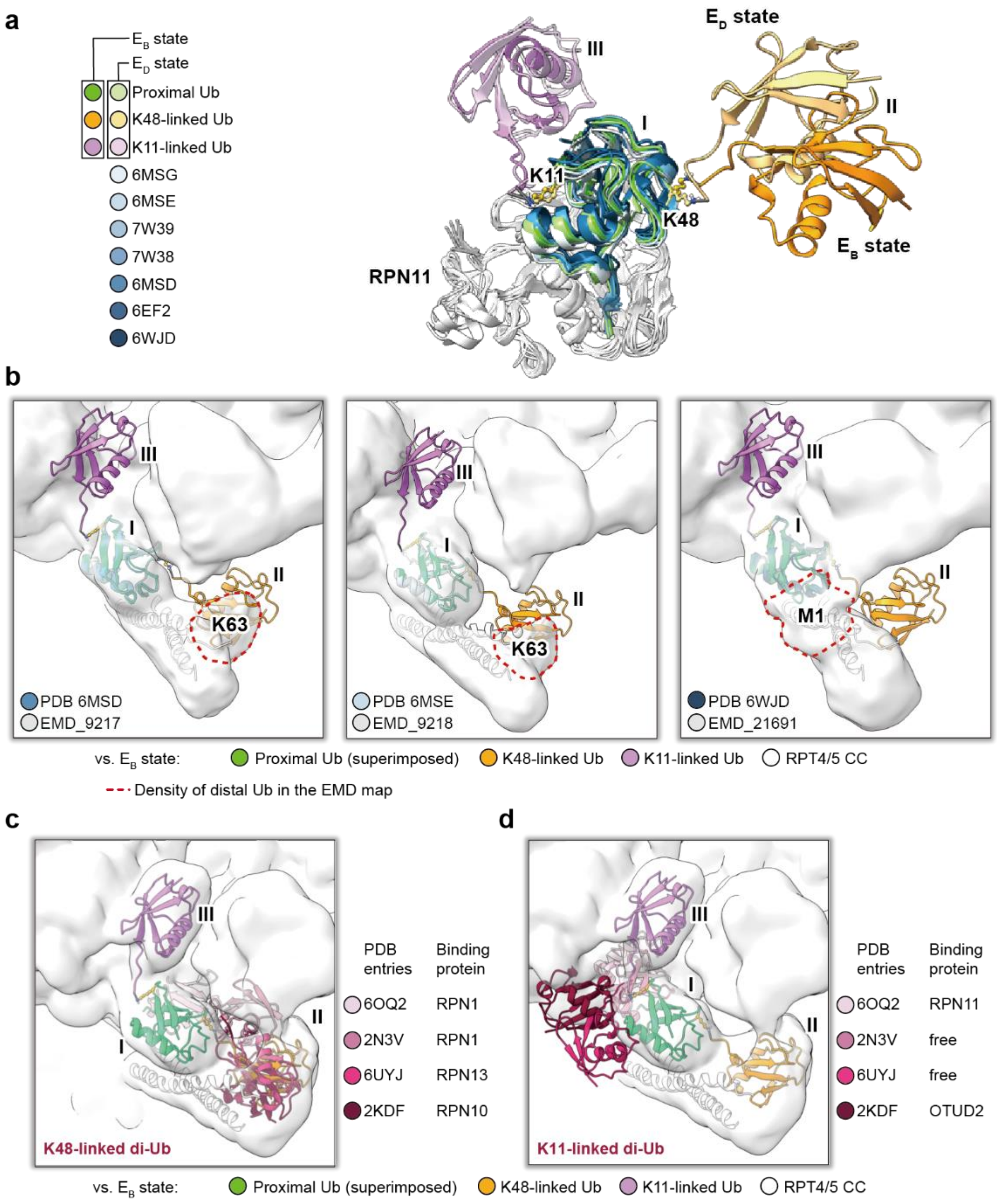

**Supplementary Fig. 7. A comparison of the proteasome-bound K11/K48-branched Ub with the available structural data. a**, Superposition of the structures of reported RPN11-bound Ub showed a highly conserved pose of the RPN11-bound proximal Ub. The individual structures are color coded as indicated on the left with their corresponding PDB ID. **b**, Superposition of reported EM maps of the Ub-bound RPT4/5 CC with partially resolved EM density of the proximal Ub and K48-linked Ub (the equivalent of $Ub_{II}$ in this study). The PDB IDs and the corresponding EMDB IDs are indicated on the lower left corner of individual panels. The positions of the observed Ub in the earlier studies are outlined in dashed red lines, and they deviate significantly from the $Ub_{II}$ in this study. **c**, Superposition of K48-linked di-Ub in complex with different Ub receptors with respect to $Ub_{I}$ resolved in this study shows a broad conformation space sampled by the distal Ubs in the previously reported structures. Nevertheless, the distal Ubs broadly sample the same conformational space as that of $Ub_{II}$ resolved in this study. **d**, Superposition of reported K11-linked di-Ub structures shows that the distal Ubs sample different conformations distinct from the K11-linked $Ub_{III}$ resolved in this study.

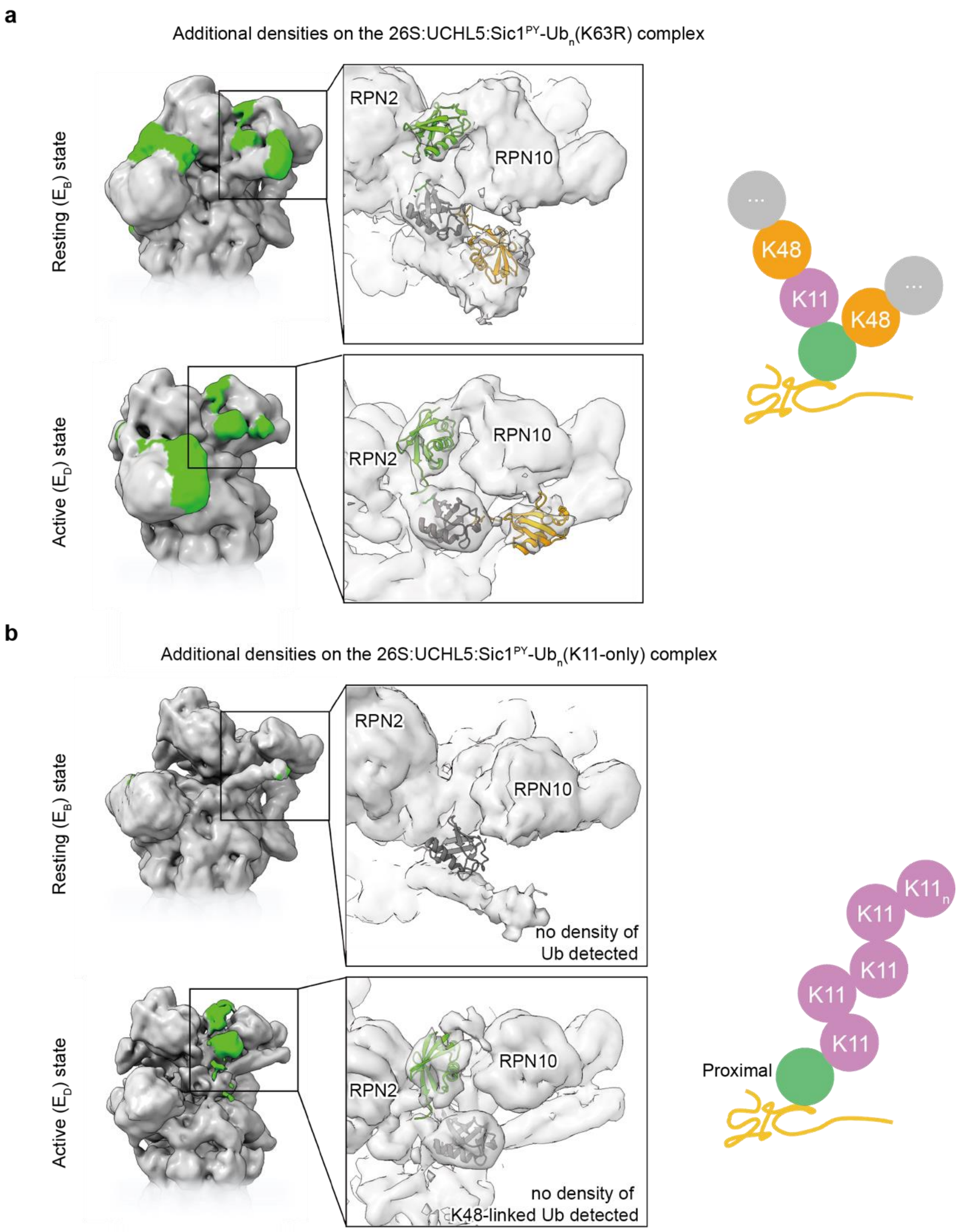

**Supplementary Fig. 8. Newly identified K11-linked Ub binding groove is specific tailored for K11-linked Ub chains.** Comparison of the cryo-EM maps of the 26S proteasome in complex with the K11/K48-branched Ub chain (**a**) and in complex with a K11-only Ub chain (**b**). The K11-only Ub chain was visible in the $E_D$ state within the K11-linked Ub binding groove (highlighted in green), but it is absent in the $E_A$ state. No EM density was was observed in the proximity of the RPT4/5 CC, which binds to K48-linked Ub extending from the proximal Ub.

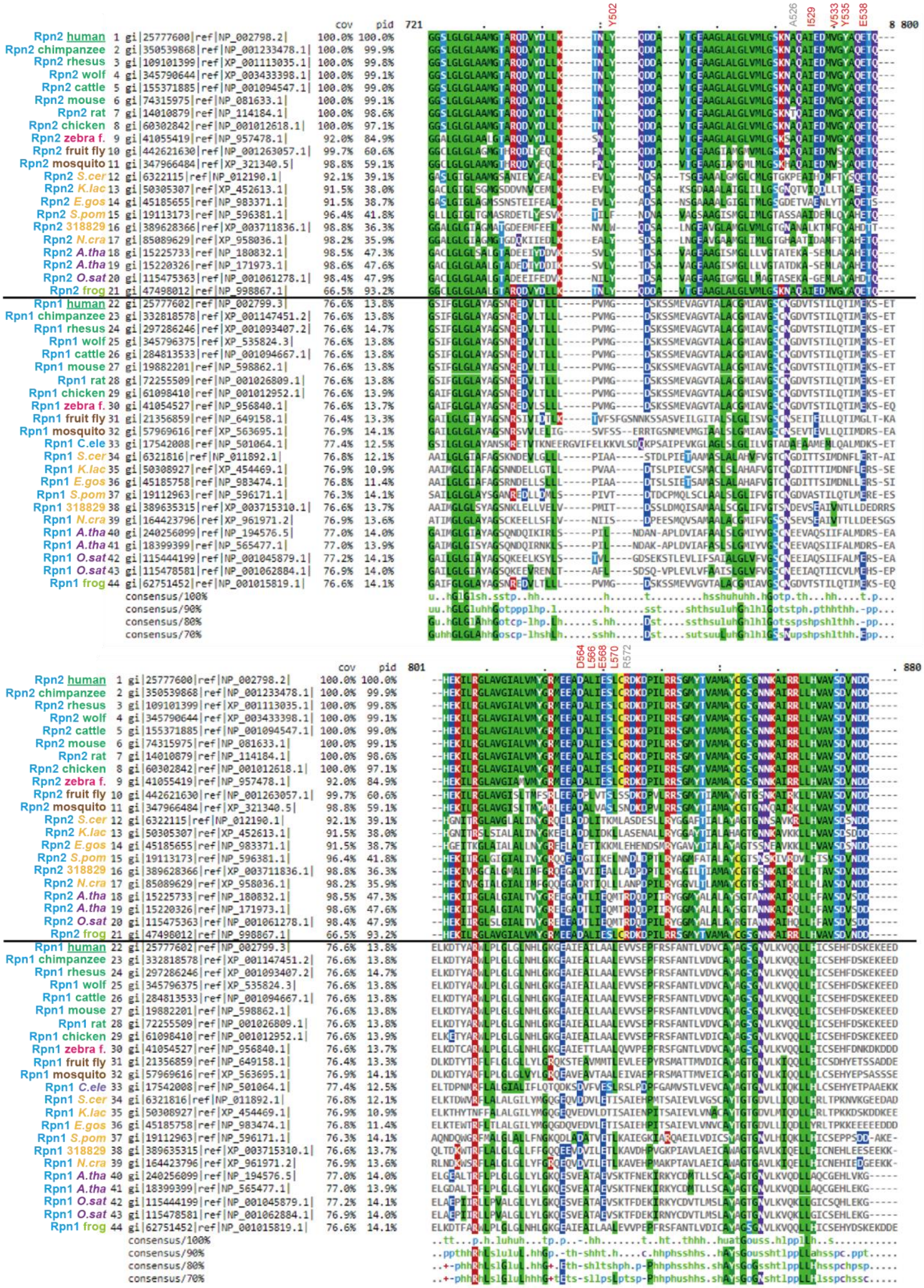

**Supplementary Fig. 9. Alignment of RPN1 and RPN2 sequences from different model species revealed a well-conserved Ub binding motif shared by the two subunits.**

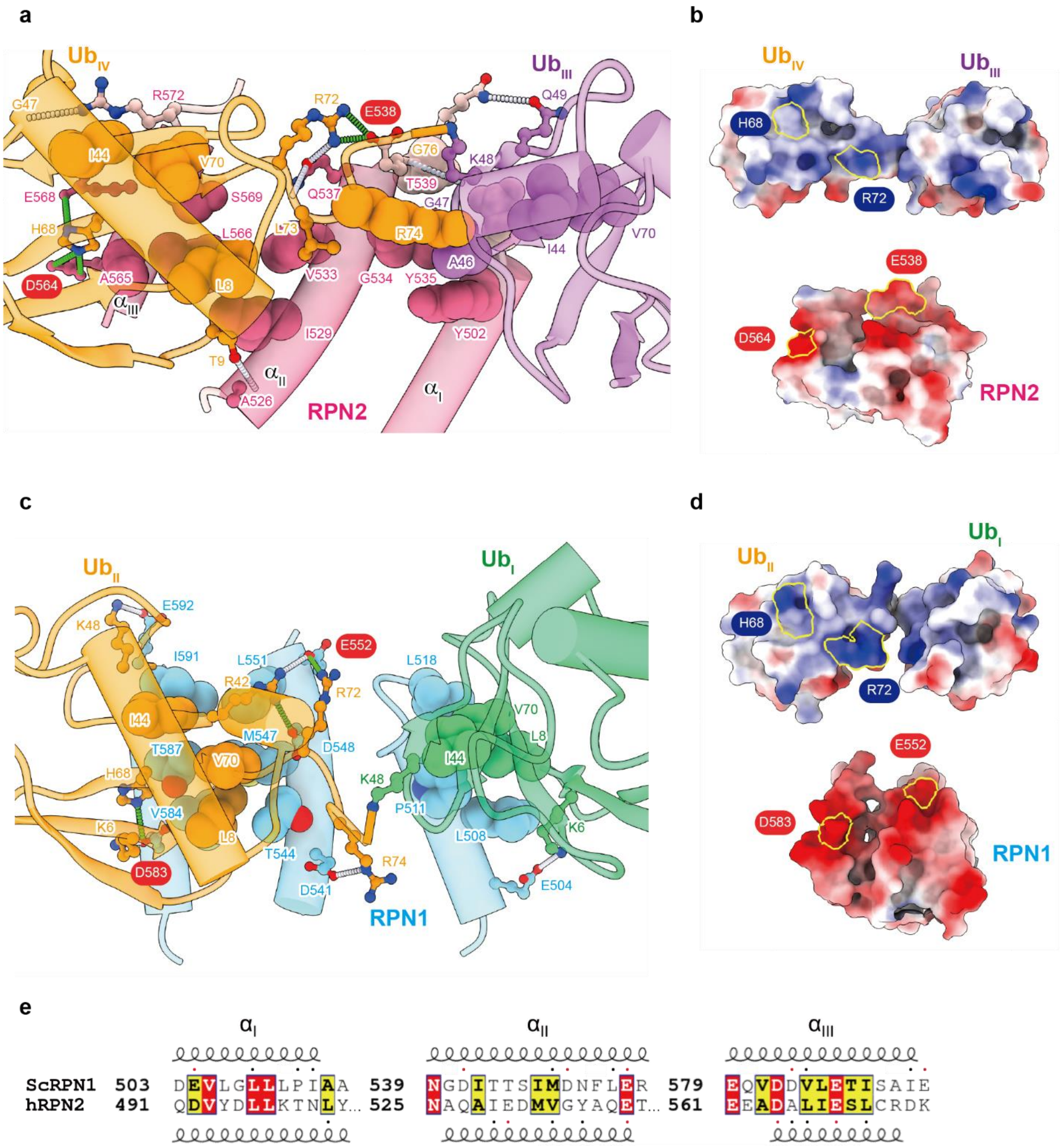

**Supplementary Fig. 10. Structural insights into the conservation of the K48-link di-Ub recognition by RPN2 and RPN1 a**, Expanded view of K48-linked $Ub_{III}$-$Ub_{IV}$ in complex with RPN2 in the $E_D$:$Ub_4$ state. **b**, Open book view of the electrostatic surface representations of K48-linked Ub and RPN2, illustrating highly complementary surface electrostatic potentials of the two molecules. The locations of the residues involved in the conserved salt bridge pairs are outlined in solid yellow lines. **c**, Expanded view of K48-linked di-Ub in complex with the T1 binding site of yeast RPN1 (*Sc*RPN1; PDB ID: 2N3V). **d**, Open book view of the electrostatic surface representation of K48-linked d-Ub and *Sc*RPN1 following the same scheme as (**b**). **e**,

Sequence alignment of the K48-linked di-Ub binding motif of human RPN2 (hRPN2) and *Sc*RPN1. The structure-based alignment was made by ESPript3.0 (https://espript.ibcp.fr/ESPript/ESPript/). Identical acidic residues are boxed with a red background, which is also outlined in red in (**a**) and (**c**). Similar hydrophobic residues are boxed with a yellow background.

**a**

RMSD (Å) values between Cα atoms of aligned 19S/RP

| | apo $E_A$ | $E_A$:$Ub_3$ | $E_B$:$Ub_4$ | $E_D$:$Ub_4$ | $E_A$/$E_{A1}$ 6msb | $E_{A2}$ 6msd | $E_B$ 6mse | $E_{C1}$ 6msg | $E_{C2}$ 6msh | $E_{D1}$ 6msj | $E_{D2}$ 6msk |
|---|---|---|---|---|---|---|---|---|---|---|---|
| apo $E_A$ | | 1.3 | 5.0 | 18.0 | 3.2 | 3.3 | 10.6 | 19.0 | 18.4 | 18.3 | 17.9 |
| $E_A$:$Ub_3$ | | | 5.0 | 17.5 | 3.1 | 3.2 | 10.7 | 19.0 | 18.4 | 18.2 | 17.4 |
| $E_B$:$Ub_4$ | | | | 19.7 | 7.0 | 5.1 | 8.7 | 19.7 | 18.6 | 20.0 | 22.3 |
| $E_D$:$Ub_4$ | | | | | 17.0 | 18.3 | 20.1 | 9.3 | 8.0 | 3.7 | 3.6 |

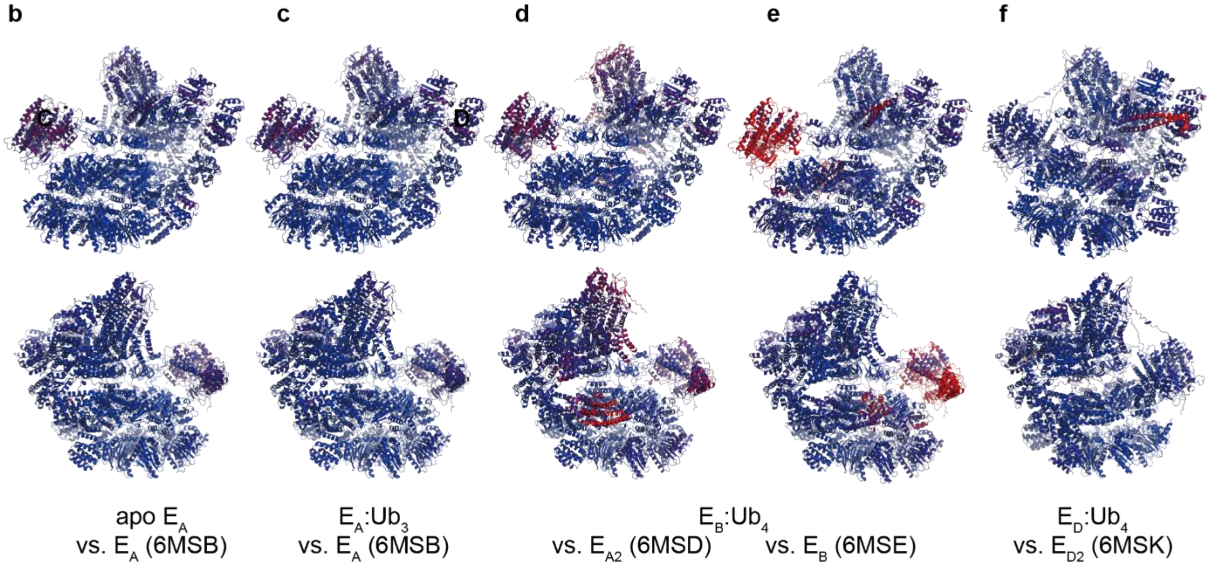

**Supplementary Fig. 11. Comparison between the proteasome states observed in the presence of K11/K48-branched ubiquitin chain and previously published states of the human 26S proteasome. a**, The Root Mean Square Deviation (RMSD) between aligned Cα within proteasomal 19S/RP of the analyzed atomic models is shown in the table. Colors varying from green to red indicate the lowest (most similar structures) and highest RMSD values, respectively. **b-f**, The 26S states reported here are colored according to the RMSD values calculated for Cα atoms of the model and the model that showed to be the most similar, as indicated by black frames in the table A. The $E_A$:$Ub_4$ state was compared to $E_{A1}$ (**d**), and $E_B$ (**e**) state, even though it showed the lowest RMSD when compared to the RP in the $E_{A1}$ state. Comparing panels (**d**) and (**e**), it can be seen that the $E_A$:$Ub_4$ shares some features of both of those states. While the RP base of the $E_A$:$Ub_4$ resembles the $E_{A1}$ state (PDB ID: 6MSD) (including the distribution of the nucleotides), the overall conformation of the lid of the $E_A$:$Ub_4$ RP is more similar to the $E_B$ state (PDB ID: 6MSE). The main differences between the $E_A$:$Ub_4$ and the earlier reported $E_B$ state are the conformations of the RPT2, RPT6, and RPN1. Compared to the $E_A$ state, the $E_A$:$Ub_4$ shows a shift and downward movement of the whole lid comparable to the conformation of the $E_B$ state.

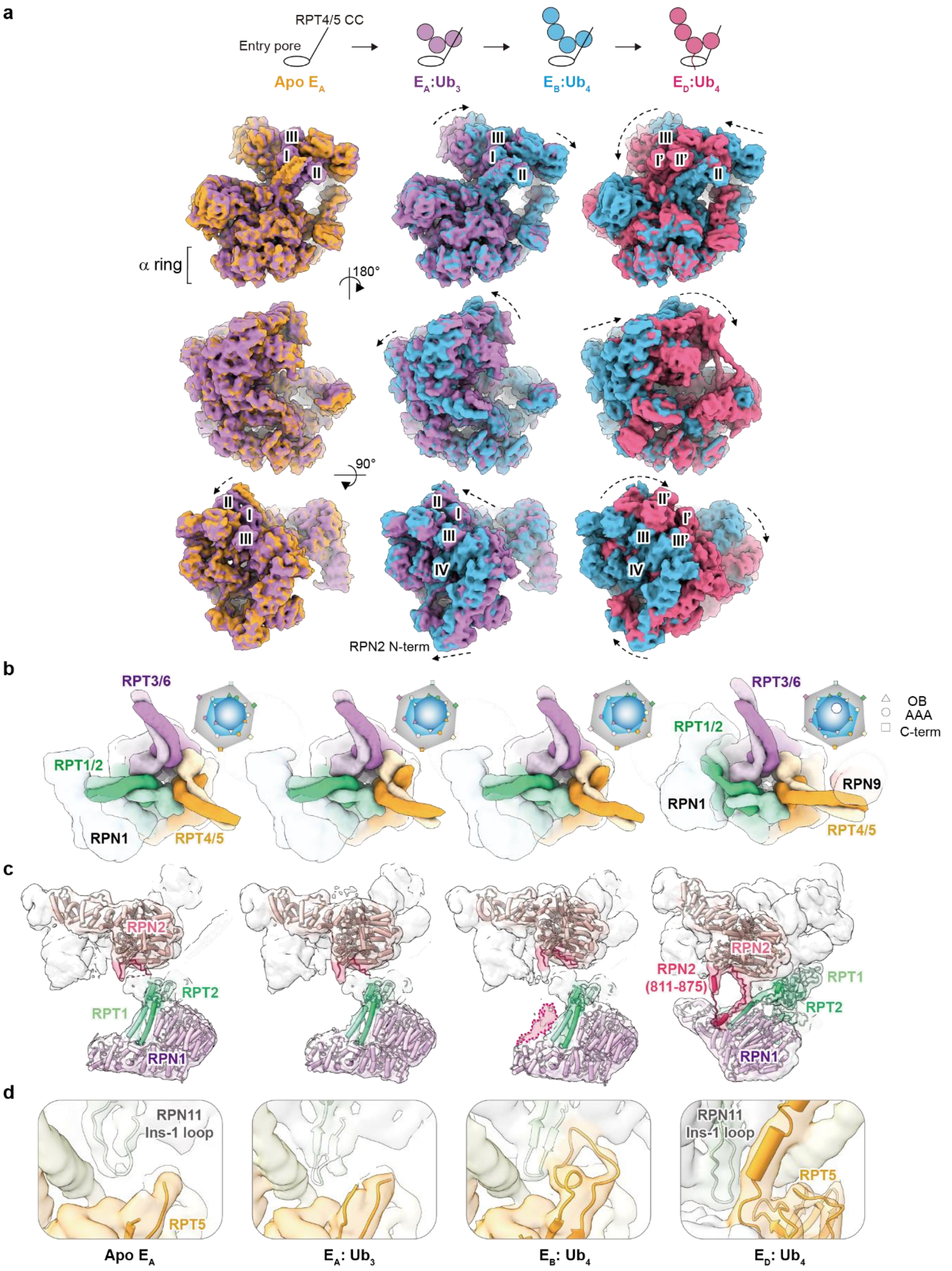

**Supplementary Fig. 12. Conformational transitions of the 19S RP induced by substrate binding. a**, Superpositions of the substrate-free $E_A$ (orange), $E_A$:$Ub_3$ (purple), $E_B$:$Ub_4$ (light blue), and $E_D$:$Ub_4$ complex (hot pink) with the positions of the four Ubs indicated in Roman numbers. **b**, Conformational transitions of the three CCs around the base of the 19S RP. Schematic representations of the OB, AAA, and C-terminal regions of the six subunits of the AAA+ ATPase are shown in triangles, circles, and squares, as indicated on the upper right corners. Note that the three structural motifs are coaxially aligned in the $E_D$:$Ub_4$ state, resulting in the appearance of the proteolytic chamber entrance as indicated by an open circle in the schematic drawing. **c**, Geometric analysis of the lever motion of the RPT4/5 CC between the four functional states. The most significant lever motion occurs during the transition from the $E_B$:$Ub_4$ (light blue) to the $E_D$:$Ub_4$ complex (hot pink). **d**, RPT1, RPT2, and RPT1 conformational changes accompany the structural ordering of the RPN2 811-875 loop. The intrinsically disordered RPN2 loop is resolved in the $E_D$:$Ub_4$ complex, allowing for atomic modeling as shown in the hot pink cartoon representation. **e**, Structural ordering of the RPN11 Ins-1 loop exhibits different conformations in the $E_B$:$Ub_4$ and $E_D$:$Ub_4$ complex, while it is disordered in the apo and $E_B$:$Ub_4$ complex.

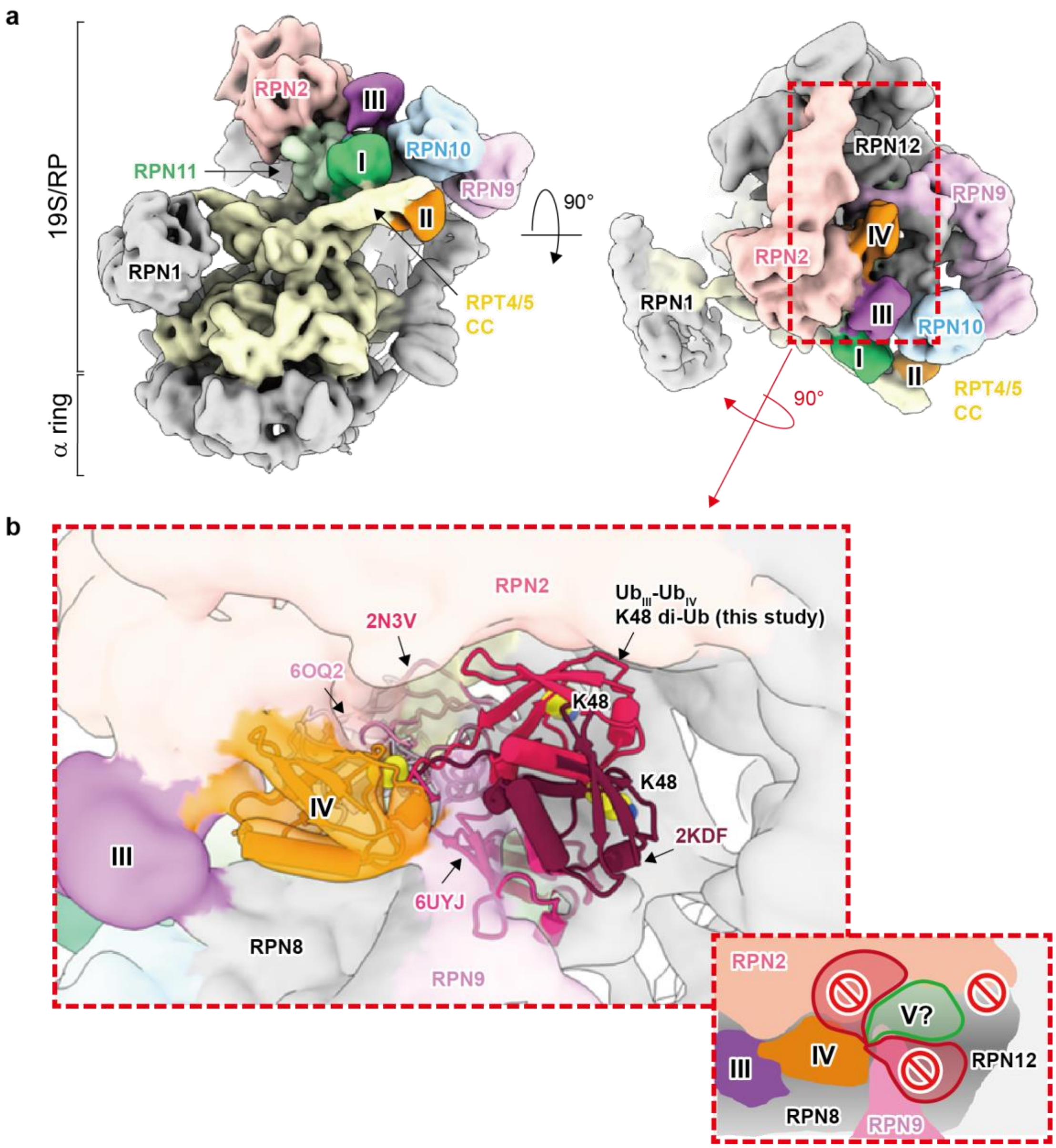

**Supplementary Fig. 13. Geometrical constrain of the K11-linked Ub binding groove for K11-linked Ub chains. a**, Orthogonal views of the cryo-EM map of the 19S RP in the $E_B$:$Ub_4$ state with the individual components colored in the same scheme as Figure 1d. **b**, Expanded view of the K11-linked Ub binding groove by RPN2, RPN8, RPN9, RPN10 and RPN11 (the last two subunits are not visible in this view). The segments of the EM map corresponding to $Ub_{III}$ and $Ub_{IV}$ are colored magenta and orange, respectively. The atomic model of $Ub_{IV}$ is shown in cartoon representation and superimposed with the structures of the previously reported K48-linked di-Ub (the PDB IDs are indicated alongside). Some of the K48-linked Ub extending from $Ub_{IV}$ will clash with the K11-Ub binding groove (such as 6UYJ, 6OQ2, and 2N3V), as schematically illustrated in the inset. However, the structure of 2KDF could still fit into the K11-linked Ub binding groove without steric clash, but further extension will clash with RPN12. The structural modeling suggests that the K11-linked Ub binding groove can accommodate up to three Ub moieties through the ubique alternating K11-K48-linkage with a further extension from $Ub_{IV}$.

**Supplementary Table 1. List of residues forming hydrogen bonds (HB, in black) and salt bridges (SB, in blue) between K11/K48-branched Ub chain and the human 26S proteasome in the observed conformational states of the complex.**

| | $E_A$:$Ub_3$ state | | | | $E_B$:$Ub_4$ state | | | | $E_D$:$Ub_4$ state | | | |
|---|---|---|---|---|---|---|---|---|---|---|---|---|
| | Ub residue (atom) | Dist. (Å) | Interacting subunit | Sub. residue (atom) | Ub residue (atom) | Dist. (Å) | Interacting subunit | Sub. residue (atom) | Ub residue (atom) | Dist. (Å) | Interacting subunit | Sub. residue (atom) |
| $Ub_I$ | GLN 40 (NE2) | 3.59 | Rpn11 | ASN 128 (OD1) | LEU 8 (O) | 3.72 | Rpn11 | VAL 108 (N) | LEU 8 (O) | 3.49 | Rpn11 | VAL 108 (N) |
| | GLN 49 (NE2) | 3.36 | Rpn11 | ASP 88 (OD2) | LEU 73 (O) | 3.66 | Rpn11 | ALA 86 (N) | GLN 40 (OE1) | 2.80 | Rpn11 | SER 132 (OG) |
| | ARG 72 (NE) | 3.00 | Rpn11 | GLU 85 (OE2) | ARG 74 (O) | 3.22 | Rpn11 | THR 129 (OG1) | LEU 73 (O) | 3.82 | Rpn11 | ALA 86 (N) |
| | ARG 72 (NH2) | 3.13 | Rpn11 | GLU 85 (OE1) | GLY 75 (O) | 3.22 | Rpn11 | VAL 84 (N) | ARG 74 (O) | 2.73 | Rpn11 | THR 129 (OG1) |
| | ARG 74 (N) | 3.78 | Rpn11 | THR 129 (OG1) | ARG 74 (NH2) | 2.43 | Rpn11 | GLU 85 (OE2) | GLY 75 (O) | 2.93 | Rpn11 | VAL 84 (N) |
| | ARG 74 (NE) | 2.57 | Rpn11 | GLU 85 (OE1) | GLY 75 (N) | 3.40 | Rpn11 | VAL 84 (O) | GLN 40 (NE2) | 3.69 | Rpn11 | ASN 128 (O) |
| | GLY 75 (N) | 2.75 | Rpn11 | VAL 84 (O) | GLY 76 (N) | 3.09 | Rpn11 | ASP 126 (OD1) | GLN 40 (NE2) | 3.52 | Rpn11 | ASN 128 (OD1) |
| | GLY 76 (N) | 3.59 | Rpn11 | GLU 52 (OE2) | ARG 74 (NE) | 2.89 | Rpn11 | GLU 85 (OE2) | ARG 74 (N) | 3.37 | Rpn11 | THR 129 (OG1) |
| | GLY 76 (N) | 2.64 | Rpn11 | SER 83 (OG) | ARG 74 (NH2) | 2.43 | Rpn11 | GLU 85 (OE2) | ARG 74 (NH1) | 2.77 | Rpn11 | GLU 85 (OE2) |
| | GLY 76 (N) | 3.19 | Rpn11 | ASP 126 (OD1) | ASP 58 (OD2) | 2.43 | Rpt5 | ARG 61 (NH2) | GLY 75 (N) | 3.12 | Rpn11 | VAL 84 (O) |
| | LEU 8 (O) | 3.66 | Rpn11 | VAL 108 (N) | ASP 58 (OD2) | 3.89 | Rpt5 | ARG 61 (NE) | GLY 76 (N) | 3.52 | Rpn11 | ASP 126 (OD1) |
| | LEU 73 (O) | 2.76 | Rpn11 | ALA 86 (N) | ASP 58 (OD2) | 2.43 | Rpt5 | ARG 61 (NH2) | ARG 74 (NE) | 4.00 | Rpn11 | GLU 85 (OE2) |
| | ARG 74 (O) | 2.39 | Rpn11 | THR 129 (OG1) | | | | | ARG 74 (NH1) | 2.77 | Rpn11 | GLU 85 (OE2) |
| | GLY 76 (OXT) | 2.72 | Rpn11 | SER 123 (OG) | | | | | | | | |
| | ARG 72 (NE) | 3.00 | Rpn11 | GLU 85 (OE2) | | | | | | | | |
| | ARG 72 (NE) | 3.83 | Rpn11 | GLU 85 (OE1) | | | | | | | | |
| | ARG 72 (NH2) | 3.58 | Rpn11 | GLU 85 (OE2) | | | | | | | | |
| | ARG 72 (NH2) | 3.13 | Rpn11 | GLU 85 (OE1) | | | | | | | | |
| | ARG 74 (NE) | 2.57 | Rpn11 | GLU 85 (OE1) | | | | | | | | |
| | ARG 74 (NH2) | 2.72 | Rpn11 | GLU 85 (OE1) | | | | | | | | |
| | ASP 58 (OD2) | 3.08 | Rpt5 | ARG 61 (NH2) | | | | | | | | |
| | ASP 58 (OD2) | 3.08 | Rpt5 | ARG 61 (NH2) | | | | | | | | |
| $Ub_{II}$ | GLU 34 (O) | 2.91 | Rpt4 | ARG 30 (NH1) | THR 9 (OG1) | 3.38 | Rpt4 | LYS 20 (NZ) | THR 22 (OG1) | 3.50 | Rpt4 | ARG 25 (NH1) |
| | GLU 34 (OE1) | 3.98 | Rpt4 | LYS 27 (NZ) | GLU 34 (OE2) | 2.66 | Rpt4 | ARG 30 (NH2) | THR 22 (OG1) | 3.54 | Rpt4 | ARG 25 (NH2) |
| | GLN 40 (NE2) | 3.32 | Rpt5 | GLU 58 (OE2) | GLU 34 (OE2) | 2.23 | Rpt4 | LYS 27 (NZ) | ASP 52 (OD2) | 2.42 | Rpt5 | THR 63 (OG1) |
| | | | | | GLU 34 (OE1) | 3.77 | Rpt4 | LYS 27 (NZ) | | | | |
| | | | | | GLU 34 (OE2) | 3.33 | Rpt4 | ARG 30 (NH1) | | | | |
| | | | | | GLU 34 (OE2) | 2.66 | Rpt4 | ARG 30 (NH2) | | | | |
| | | | | | GLU 34 (OE2) | 2.23 | Rpt4 | LYS 27 (NZ) | | | | |
| | | | | | THR 9 (OG1) | 2.55 | Rpt5 | GLU 51 (OE2) | | | | |
| | | | | | ARG 74 (NH1) | 2.91 | Rpt5 | GLU 58 (OE2) | | | | |
| | | | | | ARG 74 (NH2) | 3.28 | Rpt5 | GLU 58 (OE2) | | | | |
| | | | | | ARG 74 (NH1) | 2.91 | Rpt5 | GLU 58 (OE2) | | | | |
| $Ub_{III}$ | ARG 74 (NH1) | 3.02 | Rpn11 | PHE 61 (O) | ARG 72 (NH1) | 3.84 | Rpn11 | GLU 60 (OE1) | ARG 42 (NH2) | 3.08 | Rpn11 | GLU 60 (OE2) |
| | ARG 74 (NH2) | 3.12 | Rpn11 | PHE 61 (O) | ARG 72 (NH2) | 3.53 | Rpn11 | GLU 60 (OE1) | ARG 42 (NH2) | 3.08 | Rpn11 | GLU 60 (OE2) |
| | ARG 54 (NH2) | 2.75 | Rpn8 | ASN 31 (OD1) | ARG 72 (NH1) | 3.84 | Rpn11 | GLU 60 (OE1) | GLY 53 (O) | 3.57 | Rpn8 | ASN 31 (N) |
| | GLU 51 (OE1) | 2.96 | Rpn8 | LYS 28 (NZ) | ARG 72 (NH2) | 3.53 | Rpn11 | GLU 60 (OE1) | | | | |
| | GLU 51 (OE2) | 3.72 | Rpn8 | LYS 28 (NZ) | GLU 51 (OE2) | 3.16 | Rpn8 | LYS 28 (NZ) | | | | |
| | GLU 51 (OE1) | 2.96 | Rpn8 | LYS 28 (NZ) | GLU 51 (OE2) | 3.16 | Rpn8 | LYS 28 (NZ) | | | | |
| | | | | | GLN 49 (OE1) | 3.69 | Rpn2 | GLN 540 (NE2) | | | | |
| | | | | | LYS 48 (N) | 3.79 | Rpn2 | GLU 538 (O) | | | | |
| $Ub_{IV}$ | | | | | GLY 47 (O) | 3.39 | Rpn2 | ARG 572 (NH2) | GLY 47 (O) | 3.43 | Rpn2 | ARG 572 (NH2) |
| | | | | | THR 9 (OG1) | 2.30 | Rpn2 | ALA 526 (O) | THR 9 (OG1) | 3.38 | Rpn2 | ALA 526 (O) |
| | | | | | LEU 73 (N) | 3.46 | Rpn2 | GLN 537 (OE1) | LEU 73 (N) | 3.65 | Rpn2 | GLN 537 (OE1) |
| | | | | | ARG 72 (NH1) | 2.55 | Rpn2 | GLN 537 (OE1) | ARG 74 (NE) | 2.37 | Rpn2 | GLU 538 (OE2) |
| | | | | | ARG 72 (NH1) | 2.57 | Rpn2 | GLU 538 (OE2) | LYS 6 (NZ) | 3.61 | Rpn2 | ASP 564 (OD1) |
| | | | | | ARG 72 (NH2) | 2.67 | Rpn2 | GLU 538 (OE2) | ARG 74 (NE) | 3.45 | Rpn2 | GLU 538 (OE1) |
| | | | | | HIS 68 (NE2) | 3.22 | Rpn2 | ASP 564 (OD1) | ARG 74 (NE) | 2.37 | Rpn2 | GLU 538 (OE2) |
| | | | | | ARG 72 (NH1) | 2.57 | Rpn2 | GLU 538 (OE2) | ARG 74 (NH2) | 2.65 | Rpn2 | GLU 538 (OE2) |
| | | | | | ARG 72 (NH2) | 2.67 | Rpn2 | GLU 538 (OE2) | LYS 6 (NZ) | 3.61 | Rpn2 | ASP 564 (OD1) |
| | | | | | HIS 68 (NE2) | 3.22 | Rpn2 | ASP 564 (OD1) | HIS 68 (ND1) | 3.31 | Rpn2 | GLU 568 (OE2) |
| | | | | | HIS 68 (NE2) | 3.95 | Rpn2 | ASP 564 (OD2) | HIS 68 (NE2) | 2.84 | Rpn2 | GLU 568 (OE2) |
| | | | | | HIS 68 (ND1) | 3.83 | Rpn2 | GLU 568 (OE2) | ASP 58 (OD2) | 3.41 | Rpn9 | LYS 361 (NZ) |
| | | | | | ASP 58 (OD2) | 2.95 | Rpn9 | LYS 361 (NZ) | ARG 54 (NH1) | 3.09 | Rpn9 | GLU 368 (OE1) |
| | | | | | ARG 54 (NH1) | 3.47 | Rpn9 | GLU 368 (OE2) | ARG 54 (NH2) | 3.80 | Rpn9 | GLU 368 (OE2) |
| | | | | | ARG 54 (NH2) | 2.44 | Rpn9 | GLU 368 (OE1) | ASP 58 (OD2) | 3.41 | Rpn9 | LYS 361 (NZ) |
| | | | | | ASP 58 (OD2) | 2.95 | Rpn9 | LYS 361 (NZ) | ARG 54 (NH1) | 3.09 | Rpn9 | GLU 368 (OE1) |
| | | | | | ARG 54 (NH1) | 3.85 | Rpn9 | GLU 368 (OE1) | ARG 54 (NH2) | 3.53 | Rpn9 | GLU 368 (OE1) |
| | | | | | ARG 54 (NH1) | 3.47 | Rpn9 | GLU 368 (OE2) | ARG 54 (NH2) | 3.80 | Rpn9 | GLU 368 (OE2) |
| | | | | | ARG 54 (NH2) | 3.99 | Rpn9 | GLU 364 (OE2) | | | | |
| | | | | | ARG 54 (NH2) | 2.44 | Rpn9 | GLU 368 (OE1) | | | | |
| | | | | | ARG 54 (NH2) | 3.28 | Rpn9 | GLU 368 (OE2) | | | | |

**Supplementary Table 2. List of residues at the interface between K11/K48-branched Ub chain and the human 26S proteasome with their solvation energy change ($\Delta^iG$ in kcal/mol) upon complex formation. Energetically favorable changes are highlighted in green.**

| | $E_A$:$Ub_3$ state | | | | | $E_B$:$Ub_4$ state | | | | | $E_D$:$Ub_4$ state | | | | |
|---|---|---|---|---|---|---|---|---|---|---|---|---|---|---|---|
| | Ub residue | $\Delta^iG$ | Interacting subunit | Sub. residue | $\Delta^iG$ | Ub residue | $\Delta^iG$ | Interacting subunit | Sub. residue | $\Delta^iG$ | Ub residue | $\Delta^iG$ | Interacting subunit | Sub. residue | $\Delta^iG$ |
| $Ub_I$ | LEU 73 | 1.8 | Rpn11 | MET 107 | 1.5 | LEU 73 | 1.8 | Rpn11 | LEU 136 | 1.62 | LEU 73 | 1.91 | Rpn11 | LEU 136 | 1.56 |
| | LEU 8 | 1.08 | Rpn11 | LEU 136 | 1.5 | LEU 8 | 1.19 | Rpn11 | MET 107 | 1.51 | LEU 8 | 1.06 | Rpn11 | MET 107 | 1.48 |
| | LEU 71 | 1.05 | Rpn11 | PRO 89 | 1.23 | ILE 44 | 0.84 | Rpn11 | PRO 89 | 1.23 | THR 9 | 1.03 | Rpn11 | PRO 89 | 1.27 |
| | ILE 44 | 0.85 | Rpn11 | VAL 125 | 0.81 | ILE 36 | 0.82 | Rpn11 | VAL 90 | 1.12 | LEU 71 | 0.95 | Rpn11 | VAL 125 | 0.98 |
| | THR 9 | 0.84 | Rpn11 | VAL 90 | 0.75 | LEU 71 | 0.82 | Rpn11 | THR 129 | 0.96 | ILE 44 | 0.93 | Rpn11 | VAL 90 | 0.87 |
| | ILE 36 | 0.83 | Rpn11 | MET 54 | 0.7 | VAL 70 | 0.71 | Rpn11 | VAL 125 | 0.87 | VAL 70 | 0.72 | Rpn11 | ALA 93 | 0.82 |
| | VAL 70 | 0.65 | Rpn11 | ALA 93 | 0.66 | GLY 76 | 0.71 | Rpn11 | ALA 93 | 0.86 | LYS 11 | 0.51 | Rpn11 | HIS 113 | 0.72 |
| | GLY 75 | 0.36 | Rpn11 | PHE 133 | 0.6 | GLY 10 | 0.32 | Rpn11 | MET 54 | 0.83 | ILE 36 | 0.42 | Rpn11 | MET 54 | 0.7 |
| | GLY 10 | 0.32 | Rpn11 | VAL 108 | 0.5 | GLY 75 | 0.18 | Rpn11 | PHE 133 | 0.61 | ARG 42 | 0.28 | Rpn11 | PHE 133 | 0.65 |
| | ARG 42 | 0.2 | Rpn11 | SER 132 | 0.49 | THR 9 | 0.18 | Rpn11 | SER 132 | 0.56 | GLY 75 | 0.28 | Rpn11 | VAL 108 | 0.59 |
| | LEU 69 | 0.17 | Rpn11 | LEU 96 | 0.38 | LYS 11 | 0.15 | Rpn11 | ALA 135 | 0.35 | GLY 10 | 0.23 | Rpn11 | HIS 115 | 0.47 |
| | THR 7 | 0.15 | Rpn11 | ALA 86 | 0.38 | ARG 42 | 0.14 | Rpn11 | VAL 108 | 0.34 | THR 7 | 0.23 | Rpn11 | LEU 96 | 0.34 |
| | HIS 68 | 0.12 | Rpn11 | SER 137 | 0.31 | LEU 69 | 0.13 | Rpn11 | MET 75 | 0.31 | HIS 68 | 0.18 | Rpn11 | SER 137 | 0.34 |
| | PRO 37 | 0.04 | Rpn11 | SER 83 | 0.22 | THR 7 | 0.1 | Rpn11 | LEU 96 | 0.29 | LEU 69 | 0.06 | Rpn11 | ALA 135 | 0.31 |
| | ARG 74 | 0.02 | Rpn11 | VAL 82 | 0.15 | LYS 48 | 0.06 | Rpn11 | SER 83 | 0.29 | GLY 47 | 0.03 | Rpn11 | MET 75 | 0.28 |
| | GLU 34 | -0.05 | Rpn11 | SER 123 | 0.14 | PRO 37 | 0.06 | Rpn11 | SER 137 | 0.26 | PRO 37 | 0 | Rpn11 | SER 83 | 0.28 |
| | GLY 47 | -0.07 | Rpn11 | MET 75 | 0.14 | GLY 35 | 0.05 | Rpn11 | LEU 56 | 0.24 | GLY 35 | -0.15 | Rpn11 | SER 132 | 0.24 |
| | ARG 72 | -0.19 | Rpn11 | LEU 56 | 0.13 | GLY 47 | 0.04 | Rpn11 | TRP 111 | 0.24 | ARG 72 | -0.19 | Rpn11 | ALA 86 | 0.22 |
| | GLN 49 | -0.24 | Rpn11 | THR 129 | 0.12 | ILE 13 | 0 | Rpn11 | ALA 86 | 0.11 | GLY 76 | -0.2 | Rpn11 | TRP 111 | 0.19 |
| | GLY 76 | -0.27 | Rpn11 | TRP 111 | 0.12 | LEU 43 | 0 | Rpn11 | HIS 113 | 0.09 | GLN 49 | -0.3 | Rpn11 | LEU 56 | 0.19 |
| | LYS 11 | -0.29 | Rpn11 | ALA 135 | 0.06 | GLU 34 | -0.09 | Rpn11 | ASP 97 | 0.04 | GLU 34 | -0.3 | Rpn11 | SER 123 | 0.14 |
| | GLN 40 | -0.69 | Rpn11 | GLY 55 | 0.03 | GLN 49 | -0.12 | Rpn11 | LYS 94 | 0.02 | GLN 40 | -0.67 | Rpn11 | THR 129 | 0.04 |
| | | | Rpn11 | HIS 115 | 0.01 | ARG 72 | -0.18 | Rpn11 | GLY 55 | 0.02 | ARG 74 | -0.74 | Rpn11 | GLU 85 | 0.01 |
| | | | Rpn11 | HIS 113 | 0.01 | GLN 40 | -0.66 | Rpn11 | ASP 88 | 0.01 | | | Rpn11 | GLY 55 | 0.01 |
| | | | Rpn11 | GLN 131 | -0.01 | HIS 68 | -0.93 | Rpn11 | HIS 115 | -0.01 | | | Rpn11 | PHE 73 | 0.01 |
| | | | Rpn11 | GLU 138 | -0.05 | ARG 74 | -1.09 | Rpn11 | VAL 87 | -0.02 | | | Rpn11 | VAL 109 | 0 |
| | | | Rpn11 | THR 80 | -0.05 | | | Rpn11 | ASP 126 | -0.08 | | | Rpn11 | ASP 126 | -0.05 |
| | | | Rpn11 | ASP 126 | -0.06 | | | Rpn11 | VAL 84 | -0.08 | | | Rpn11 | VAL 82 | -0.14 |
| | | | Rpn11 | GLU 52 | -0.12 | | | Rpn11 | SER 123 | -0.08 | | | Rpn11 | ASP 88 | -0.14 |
| | | | Rpn11 | VAL 84 | -0.14 | | | Rpn11 | VAL 82 | -0.1 | | | Rpn11 | ASP 97 | -0.15 |
| | | | Rpn11 | ASP 88 | -0.26 | | | Rpn11 | GLU 52 | -0.13 | | | Rpn11 | GLU 52 | -0.2 |
| | | | Rpn11 | GLU 85 | -0.31 | | | Rpn11 | GLN 92 | -0.23 | | | Rpn11 | VAL 84 | -0.2 |
| | | | Rpn11 | GLN 92 | -0.33 | | | Rpn11 | ASN 128 | -0.33 | | | Rpn11 | GLN 92 | -0.25 |
| | | | Rpn11 | ASN 128 | -0.34 | | | Rpn11 | GLU 85 | -0.4 | | | Rpn11 | ASN 128 | -0.42 |
| | ARG 74 | -0.19 | Rpt4 | GLN 51 | 0.10 | | | | | | | | | | |
| | | | | | | THR 9 | 0.39 | Rpt4 | LEU 16 | 0.01 | | | | | |
| | ASP 58 | -0.2 | Rpt5 | ARG 61 | -1.35 | GLY 35 | 0.35 | Rpt4 | ASP 23 | -0.02 | | | | | |
| | ARG 54 | 0.19 | Rpt5 | LEU 60 | 0.01 | ILE 36 | 0.06 | Rpt4 | LYS 34 | -0.24 | | | | | |
| | GLY 53 | 0.09 | Rpt5 | HIS 64 | -0.01 | LYS 33 | 0 | Rpt4 | LYS 20 | -0.45 | | | | | |
| | THR 22 | 0.02 | | | | GLU 34 | -0.51 | Rpt4 | LYS 27 | -0.65 | | | | | |
| | GLU 24 | 0.03 | | | | | | Rpt4 | ARG 30 | -1.1 | | | | | |
| | THR 55 | 0.11 | | | | | | | | | | | | | |
| | | | | | | THR 55 | 0.33 | Rpt5 | LEU 60 | 0.6 | | | | | |
| | | | | | | ARG 54 | 0.33 | Rpt5 | GLU 65 | -0.02 | | | | | |
| | | | | | | THR 22 | 0.16 | Rpt5 | HIS 64 | -0.14 | | | | | |
| | | | | | | GLY 53 | 0.14 | Rpt5 | ARG 61 | -0.89 | | | | | |
| | | | | | | ASP 58 | -0.2 | | | | | | | | |
| $Ub_{II}$ | GLY 35 | 0.18 | Rpt4 | LYS 27 | 0.05 | THR 9 | 0.39 | Rpt4 | LEU 16 | 0.01 | ILE 44 | 0.45 | Rpn10 | ILE 35 | 0.57 |
| | GLU 34 | -0.42 | Rpt4 | ARG 30 | -0.98 | GLY 35 | 0.35 | Rpt4 | ASP 23 | -0.02 | GLY 47 | 0.26 | Rpn10 | HIS 76 | 0.37 |
| | PRO 37 | 0.06 | Rpt4 | LEU 17 | 0.00 | ILE 36 | 0.06 | Rpt4 | LYS 34 | -0.24 | VAL 70 | 0.01 | Rpn10 | LEU 72 | 0.32 |
| | LYS 11 | 0.06 | Rpt4 | ASP 23 | -0.02 | LYS 33 | 0 | Rpt4 | LYS 20 | -0.45 | ARG 72 | -0.02 | Rpn10 | SER 73 | 0.21 |
| | ILE 36 | 0.07 | Rpt4 | LYS 20 | -0.13 | GLU 34 | -0.51 | Rpt4 | LYS 27 | -0.65 | TYR 59 | -0.06 | Rpn10 | THR 68 | 0.18 |
| | THR 9 | -0.02 | Rpt4 | HIS 19 | 0.08 | | | Rpt4 | ARG 30 | -1.1 | ALA 46 | -0.08 | Rpn10 | CYS 37 | 0.02 |
| | | | | | | | | | | | ARG 42 | -0.3 | Rpn10 | ASP 67 | -0.01 |
| | VAL 70 | 0.22 | Rpt5 | SER 57 | 0.01 | LEU 71 | 1.05 | Rpt5 | ILE 54 | 1.4 | GLU 51 | -0.35 | Rpn10 | ASP 31 | -0.09 |
| | THR 9 | 0.21 | Rpt5 | SER 50 | 0.04 | LEU 73 | 0.35 | Rpt5 | MET 55 | 0.15 | GLN 49 | -0.54 | Rpn10 | ARG 42 | -0.16 |
| | GLN 40 | -0.48 | Rpt5 | LYS 53 | 0.36 | PRO 37 | 0.08 | Rpt5 | SER 57 | -0.09 | ARG 54 | -0.6 | Rpn10 | HIS 38 | -0.21 |
| | LEU 71 | 0.34 | Rpt5 | GLU 58 | -0.30 | LEU 8 | 0.04 | Rpt5 | GLU 51 | -0.16 | LYS 48 | -0.9 | Rpn10 | ASN 34 | -0.64 |
| | LEU 8 | 1.12 | Rpt5 | LEU 48 | 0.82 | ARG 72 | -0.02 | Rpt5 | GLU 58 | -0.34 | | | | | |
| | LEU 73 | 1.58 | Rpt5 | GLU 51 | -0.19 | THR 9 | -0.18 | Rpt5 | ARG 61 | -0.44 | THR 22 | 0.14 | Rpt4 | GLU 21 | -0.11 |
| | ILE 36 | 0.01 | Rpt5 | ILE 54 | 1.50 | GLN 40 | -0.42 | | | | THR 55 | 0.1 | Rpt4 | ARG 25 | -1.17 |
| | PRO 37 | 0.10 | Rpt5 | LEU 47 | 0.02 | ARG 74 | -1.44 | | | | ASP 52 | -0.04 | | | |
| | ARG 72 | 0.05 | | | | | | | | | SER 20 | -0.12 | | | |
| | | | | | | | | | | | GLY 53 | 0.26 | Rpt5 | LEU 60 | 0.89 |
| | | | | | | | | | | | THR 55 | 0.23 | Rpt5 | MET 55 | 0.48 |
| | | | | | | | | | | | ARG 54 | 0.21 | Rpt5 | VAL 59 | 0.45 |
| | | | | | | | | | | | ASP 58 | -0.08 | Rpt5 | THR 63 | 0.11 |
| | | | | | | | | | | | GLU 51 | -0.08 | Rpt5 | LYS 56 | 0 |
| | | | | | | | | | | | ASP 39 | -0.31 | Rpt5 | GLN 67 | -0.19 |
| | | | | | | | | | | | ASP 52 | -0.36 | | | |

| | $E_A$:$Ub_3$ state | | | | |
|---|---|---|---|---|---|
| | Ub residue | $\Delta^{i}G$ | Interacting subunit | Sub. residue | $\Delta^{i}G$ |
| **$Ub_{III}$** | GLY 75 | 0.1 | Rpn11 | GLU 60 | 0.34 |
| | GLY 76 | -0.01 | Rpn11 | VAL 62 | 0.13 |
| | ARG 72 | -0.15 | Rpn11 | MET 107 | 0.06 |
| | ARG 42 | -0.65 | Rpn11 | PRO 105 | 0.04 |
| | ARG 74 | -0.96 | Rpn11 | GLU 106 | 0.02 |
| | | | Rpn11 | SER 137 | -0.06 |
| | | | Rpn11 | PHE 61 | -0.07 |
| | | | Rpn11 | GLU 138 | -0.15 |
| | LEU 8 | 0.44 | Rpn2 | GLU 538 | 0.36 |
| | ILE 44 | 0.35 | Rpn2 | TYR 535 | 0.34 |
| | ALA 46 | 0.33 | Rpn2 | ASP 504 | 0 |
| | LYS 48 | 0.29 | Rpn2 | GLY 534 | 0 |
| | GLY 47 | 0.24 | Rpn2 | THR 499 | -0.03 |
| | HIS 68 | 0.21 | Rpn2 | GLN 540 | -0.13 |
| | VAL 70 | 0.2 | Rpn2 | TYR 502 | -0.13 |
| | GLN 49 | 0.1 | Rpn2 | GLN 503 | -0.74 |
| | LYS 6 | -0.83 | | | |
| | GLY 53 | 0.31 | Rpn8 | GLY 30 | 0.2 |
| | THR 55 | 0.06 | Rpn8 | VAL 29 | -0.08 |
| | GLU 51 | -0.01 | Rpn8 | GLY 27 | -0.18 |
| | ASP 52 | -0.28 | Rpn8 | ASN 31 | -0.32 |
| | ARG 54 | -0.55 | Rpn8 | LYS 28 | -0.42 |
| **$Ub_{IV}$** | | | | | |

| | $E_B$:$Ub_4$ state | | | | |
|---|---|---|---|---|---|
| | Ub residue | $\Delta^{i}G$ | Interacting subunit | Sub. residue | $\Delta^{i}G$ |
| **$Ub_{III}$** | ARG 74 | -0.77 | Rpn11 | MET 107 | 0.71 |
| | ARG 72 | -1.31 | Rpn11 | VAL 62 | 0.12 |
| | | | Rpn11 | PHE 61 | -0.12 |
| | | | Rpn11 | GLU 60 | -0.21 |
| | ALA 46 | 0.58 | Rpn2 | TYR 535 | 0.44 |
| | GLY 47 | 0.53 | Rpn2 | TYR 502 | 0.43 |
| | ILE 44 | 0.48 | Rpn2 | THR 539 | 0.11 |
| | LYS 48 | 0.4 | Rpn2 | ASP 504 | 0.05 |
| | VAL 70 | 0.34 | Rpn2 | GLU 538 | 0.04 |
| | LEU 8 | 0.27 | Rpn2 | THR 499 | -0.09 |
| | LEU 69 | 0 | Rpn2 | GLN 540 | -0.32 |
| | GLN 49 | -0.11 | Rpn2 | GLN 503 | -0.64 |
| | HIS 68 | -0.11 | | | |
| | THR 55 | 0.22 | Rpn8 | ASN 31 | 0.34 |
| | GLY 53 | 0.18 | Rpn8 | GLY 27 | 0.15 |
| | ASP 52 | 0.08 | Rpn8 | ILE 26 | 0.12 |
| | ASP 58 | -0.02 | Rpn8 | VAL 29 | 0.08 |
| | GLU 51 | -0.21 | Rpn8 | GLY 30 | 0 |
| | ARG 54 | -1.29 | Rpn8 | ARG 25 | 0 |
| | | | Rpn8 | GLN 32 | -0.02 |
| | | | Rpn8 | ASN 24 | -0.04 |
| | | | Rpn8 | LYS 28 | -0.28 |
| | GLU 24 | -0.21 | Rpn10 | LYS 103 | 0.23 |
| **$Ub_{IV}$** | LEU 8 | 1.32 | Rpn2 | ALA 565 | 0.98 |
| | LEU 73 | 1.19 | Rpn2 | ILE 529 | 0.51 |
| | ILE 44 | 0.82 | Rpn2 | VAL 533 | 0.43 |
| | VAL 70 | 0.58 | Rpn2 | GLY 534 | 0.36 |
| | HIS 68 | 0.58 | Rpn2 | ALA 526 | 0.35 |
| | ARG 74 | 0.3 | Rpn2 | LEU 566 | 0.28 |
| | GLY 76 | 0.21 | Rpn2 | SER 569 | 0.27 |
| | THR 9 | 0.11 | Rpn2 | GLU 538 | 0.14 |
| | GLY 10 | 0.11 | Rpn2 | GLU 530 | 0.1 |
| | LYS 48 | 0.03 | Rpn2 | ASP 573 | 0.04 |
| | LYS 11 | 0.03 | Rpn2 | MET 556 | 0.02 |
| | GLN 49 | 0.02 | Rpn2 | LEU 570 | 0.01 |
| | THR 7 | 0.01 | Rpn2 | GLU 562 | -0.03 |
| | GLY 47 | -0.03 | Rpn2 | LYS 524 | -0.08 |
| | LEU 71 | -0.04 | Rpn2 | ASP 564 | -0.17 |
| | LYS 6 | -0.14 | Rpn2 | ARG 572 | -0.25 |
| | ARG 42 | -0.62 | Rpn2 | GLU 568 | -0.27 |
| | ARG 72 | -1.3 | Rpn2 | LYS 574 | -0.41 |
| | | | Rpn2 | GLN 537 | -0.67 |
| | ARG 54 | -0.03 | Rpn8 | LYS 129 | 0.4 |
| | ASP 39 | -0.09 | Rpn8 | LYS 199 | -0.29 |
| | GLU 51 | -0.37 | | | |
| | THR 55 | 0.12 | Rpn9 | MET 365 | 1.22 |
| | LYS 48 | 0.06 | Rpn9 | THR 376 | 0.29 |
| | ASP 58 | 0.04 | Rpn9 | GLU 364 | -0.1 |
| | GLY 53 | -0.08 | Rpn9 | GLU 368 | -0.18 |
| | GLU 51 | -0.1 | Rpn9 | HIS 372 | -0.18 |
| | ARG 54 | -1.7 | Rpn9 | LYS 361 | -0.85 |

| | $E_D$:$Ub_4$ state | | | | |
|---|---|---|---|---|---|
| | Ub residue | $\Delta^{i}G$ | Interacting subunit | Sub. residue | $\Delta^{i}G$ |
| **$Ub_{III}$** | ARG 72 | 0.05 | Rpn11 | MET 107 | 0.35 |
| | ARG 74 | 0.03 | Rpn11 | PHE 61 | 0.34 |
| | GLY 76 | -0.13 | Rpn11 | SER 137 | 0.04 |
| | ARG 42 | -0.26 | Rpn11 | LEU 136 | -0.1 |
| | | | Rpn11 | GLU 60 | -0.44 |
| | GLN 49 | 0.4 | Rpn2 | TYR 535 | 0.51 |
| | ILE 44 | 0.39 | Rpn2 | TYR 502 | 0.34 |
| | GLY 47 | 0.34 | Rpn2 | THR 539 | 0.08 |
| | ALA 46 | 0.26 | Rpn2 | GLY 534 | -0.01 |
| | HIS 68 | 0.24 | Rpn2 | THR 499 | -0.02 |
| | LEU 8 | 0.19 | Rpn2 | ASP 504 | -0.03 |
| | VAL 70 | 0.16 | Rpn2 | ASN 470 | -0.13 |
| | LYS 48 | 0.11 | Rpn2 | GLN 540 | -0.27 |
| | THR 7 | 0.02 | Rpn2 | GLU 538 | -0.29 |
| | LYS 6 | -0.24 | Rpn2 | GLN 503 | -0.67 |
| | GLY 53 | 0.2 | Rpn8 | ASN 31 | 0.32 |
| | THR 55 | 0.13 | Rpn8 | GLY 27 | 0.27 |
| | ASP 52 | 0.11 | Rpn8 | GLY 30 | 0.18 |
| | GLU 51 | -0.09 | Rpn8 | LYS 28 | 0.16 |
| | ARG 54 | -0.71 | Rpn8 | ILE 26 | 0.04 |
| | | | Rpn8 | VAL 29 | 0.02 |
| | | | Rpn8 | ARG 25 | -0.01 |
| | | | Rpn8 | GLN 32 | -0.04 |
| | | | Rpn8 | ASN 24 | -0.05 |
| **$Ub_{IV}$** | LEU 8 | 1.26 | Rpn2 | ALA 565 | 0.92 |
| | LEU 73 | 1.11 | Rpn2 | VAL 533 | 0.64 |
| | VAL 70 | 0.88 | Rpn2 | LEU 566 | 0.44 |
| | HIS 68 | 0.88 | Rpn2 | ILE 529 | 0.4 |
| | ILE 44 | 0.54 | Rpn2 | GLY 534 | 0.3 |
| | ARG 72 | 0.36 | Rpn2 | ALA 526 | 0.2 |
| | GLN 49 | 0.13 | Rpn2 | GLU 530 | 0.15 |
| | GLY 76 | 0.1 | Rpn2 | TYR 535 | 0.12 |
| | GLY 10 | 0.07 | Rpn2 | SER 569 | 0.09 |
| | THR 7 | -0.01 | Rpn2 | LEU 570 | 0.05 |
| | LEU 69 | -0.03 | Rpn2 | ASP 564 | -0.12 |
| | THR 9 | -0.05 | Rpn2 | GLU 568 | -0.3 |
| | LEU 71 | -0.05 | Rpn2 | GLN 537 | -0.59 |
| | ALA 46 | -0.06 | Rpn2 | GLU 538 | -0.65 |
| | GLY 47 | -0.12 | Rpn2 | ARG 572 | -0.84 |
| | ARG 74 | -0.56 | | | |
| | LYS 6 | -0.57 | | | |
| | GLU 51 | -0.02 | Rpn8 | LYS 199 | -0.8 |
| | ARG 54 | -0.1 | | | |
| | THR 55 | 0.07 | Rpn9 | MET 365 | 1.13 |
| | ASP 58 | -0.02 | Rpn9 | GLU 368 | -0.32 |
| | GLY 53 | -0.07 | Rpn9 | LYS 361 | -0.56 |
| | ARG 54 | -0.98 | | | |

**Supplementary Table 3. Characteristics of interaction interfaces between K11/K48-branched tetra-Ub and human 26S proteasome. Solvent Accessible Surface Area occupied by each of the Ubs.**

| $E_B$:$Ub_4$ state | | | | | | | | | | | |
|---|---|---|---|---|---|---|---|---|---|---|---|
| Ub | | | 26S subunits | | | Interface area (Å$^2$) | $\Delta^iG$ (kcal/mol) | $\Delta^iG$ P-value | $N_{HB}$ | $N_{SB}$ | % of total interface area |
| | $^iN_{at}$ | $^iN_{res}$ | | $^iN_{at}$ | $^iN_{res}$ | | | | | | |
| **$Ub_I$** | 102 | 26 | **Rpn11** | 114 | 34 | 1157.4 | -16.6 | 0.138 | 7 | 2 | 38% |
| **$Ub_I$** | 12 | 5 | **Rpt5** | 11 | 4 | 131.5 | -0.3 | 0.510 | 1 | 2 | 4% |
| **$Ub_{II}$** | 22 | 8 | **Rpt5** | 23 | 6 | 226.8 | -0.0 | 0.626 | 2 | 2 | 7% |
| **$Ub_{II}$** | 17 | 5 | **Rpt4** | 12 | 6 | 124 | 2.2 | 0.700 | 3 | 4 | 4% |
| **$Ub_{III}$** | 30 | 9 | **Rpn2** | 37 | 8 | 289.4 | -2.4 | 0.351 | 2 | 0 | 9% |
| **$Ub_{III}$** | 22 | 6 | **Rpn8** | 27 | 9 | 198.7 | 0.7 | 0.780 | 1 | 1 | 6% |
| **$Ub_{III}$** | 8 | 2 | **Rpn11** | 13 | 4 | 79.3 | 1.6 | 0.769 | 2 | 2 | 3% |
| **$Ub_{III}$** | 1 | 1 | **Rpn10** | 4 | 1 | 15.4 | -0.0 | 0.621 | 0 | 0 | 1% |
| **$Ub_{IV}$** | 65 | 18 | **Rpn2** | 66 | 19 | 640.7 | -4.8 | 0.353 | 7 | 5 | 21% |
| **$Ub_{IV}$** | 20 | 6 | **Rpn9** | 15 | 6 | 154.7 | 1.5 | 0.775 | 3 | 6 | 5% |
| **$Ub_{IV}$** | 9 | 3 | **Rpn8** | 7 | 2 | 59.2 | 0.4 | 0.715 | 1 | 1 | 2% |
| | | | Total interface area (Å$^2$)= | | | **3077.1** | % of total interface area occupied by each Ub (Å$^2$) | | | **$Ub_I$** | 42% |
| | | | | | | | | | | **$Ub_{II}$** | 11% |
| | | | | | | | | | | **$Ub_{III}$** | 19% |
| | | | | | | | | | | **$Ub_{IV}$** | 28% |

| $E_D$:$Ub_4$ state | | | | | | | | | | | |
|---|---|---|---|---|---|---|---|---|---|---|---|
| Ub | | | 26S subunits | | | Interface area (Å$^2$) | $\Delta^iG$ (kcal/mol) | $\Delta^iG$ P-value | $N_{HB}$ | $N_{SB}$ | % of total interface area |
| | $^iN_{at}$ | $^iN_{res}$ | | $^iN_{at}$ | $^iN_{res}$ | | | | | | |
| **$Ub_I$** | 89 | 23 | **Rpn11** | 115 | 34 | 1117.2 | -17.4 | 0.099 | 11 | 2 | 38% |
| **$Ub_{II}$** | 25 | 11 | **Rpn10** | 28 | 11 | 254.3 | 1.6 | 0.759 | 3 | 2 | 9% |
| **$Ub_{II}$** | 23 | 7 | **Rpt5** | 16 | 6 | 210 | -1.6 | 0.393 | 1 | 0 | 7% |
| **$Ub_{II}$** | 8 | 4 | **Rpt4** | 6 | 2 | 57.8 | 1.2 | 0.742 | 2 | 0 | 2% |
| **$Ub_{III}$** | 36 | 10 | **Rpn2** | 40 | 10 | 304.2 | -1.4 | 0.570 | 0 | 0 | 10% |
| **$Ub_{III}$** | 18 | 5 | **Rpn8** | 23 | 9 | 168.8 | -0.5 | 0.701 | 1 | 0 | 6% |
| **$Ub_{III}$** | 12 | 4 | **Rpn11** | 12 | 5 | 90.3 | 0.1 | 0.743 | 1 | 1 | 3% |
| **$Ub_{IV}$** | 63 | 17 | **Rpn2** | 61 | 15 | 612 | -4.7 | 0.409 | 5 | 6 | 21% |
| **$Ub_{IV}$** | 14 | 4 | **Rpn9** | 7 | 3 | 103.3 | 0.8 | 0.710 | 3 | 4 | 4% |
| **$Ub_{IV}$** | 6 | 2 | **Rpn8** | 2 | 1 | 20.3 | 0.9 | 0.834 | 0 | 0 | 1% |
| | | | Total interface area (Å$^2$)= | | | **2938.2** | % of total interface area occupied by each Ub (Å$^2$) | | | **$Ub_I$** | 38% |
| | | | | | | | | | | **$Ub_{II}$** | 18% |
| | | | | | | | | | | **$Ub_{III}$** | 19% |
| | | | | | | | | | | **$Ub_{IV}$** | 25% |

$^iN_{at}$ indicates the number of interfacing atoms in the corresponding structure. $^iN_{res}$ indicates the number of interfacing residues in the corresponding structure. Interface area in Å$^2$, calculated as difference in total accessible surface areas of isolated and interfacing structures divided by two. $\Delta^iG$ indicates the solvation free energy gain upon formation of the interface. $\Delta^iG$ P-value indicates the P-value of the observed solvation free energy gain (P=0.5 means that the interface is not "surprising" at all with ΔiG equal to an average value for given structures; P>0.5 means that the interface is less hydrophobic then it could be, therefore the interface is likely to be an artefact of crystal packing; P<0.5 indicates interfaces with surprising (higher than would-be-average for given structures) hydrophobicity). $N_{HB}$ indicates the number of potential hydrogen bonds across the interface. $N_{SB}$ indicates the number of potential salt bridges across the interface.

**Supplementary Table 4. Summary of the cryo-EM data collection parameters and model statistics.**

| | Apo $E_A$<br>PDB 8JRI<br>EMD- 36598 | $E_A$:$Ub_3$<br>PDB 8JRT<br>EMD- 36605<br>EMD-37269,<br>EMD-37273,<br>EMD-37276,<br>EMD-37277 | $E_B$:$Ub_4$<br>PDB 8JTI<br>EMD-36645<br>EMD-37317<br>EMD-37319<br>EMD-37327<br>EMD-37328 | $E_D$:$Ub_4$<br>PDB 8K0G<br>EMD-36764<br>EMD-37334<br>EMD-37335<br>EMD-37341<br>EMD-37344 |
|---|---|---|---|---|
| **Data collection and processing** | | | | |
| Magnification | 64,000x | | | |
| Voltage (kV) | 300 | | | |
| Electron exposure ($e^-/Å^2$) | 48–50 | | | |
| Defocus range (μm) | -1.2 to -1.8 | | | |
| Pixel size (Å) | 0.7 super-resolution pixel | | | |
| Symmetry imposed | C1 | C1 | C1 | C1 |
| Initial particle images (no.) | 2 695 911 | | | |
| Final particle images (no.) | 153 646 | 54 794 | 52 216 | 84 476 |
| Map resolution (Å) | 3.4 | 3.6 | 3.8 | 3.8 |
| FSC threshold | 0.143 | 0.143 | 0.143 | 0.143 |
| **Refinement** | | | | |
| Initial model used (PDB ID) | 6MSB | 6MSB | 6MSE | 6MSK |
| Model resolution (Å) | 3.6 | 3.6 | 3.8 | 3.8 |
| FSC threshold | 0.143 | 0.143 | 0.143 | 0.143 |
| Model composition | | | | |
| Non-hydrogen atoms | 62 679 | 66 765 | 67 839 | 68 848 |
| Protein atoms | 8 787 | 9 029 | 9 115 | 9 220 |
| Ligands | MG:5<br>ATP:4<br>ADP:2 | MG:5<br>ATP:4<br>ADP:2 | MG:5<br>ATP:4<br>ADP:2 | MG:4<br>ATP:3<br>ADP:2 |
| *B* factors ($Å^2$) | | | | |
| Protein | 80.24 | 113.61 | 99.06 | 107.54 |
| Ligand | 31.10 | 58.82 | 47.71 | 64.33 |
| R.m.s. deviations | | | | |
| Bond lengths (Å) | 0.003 | 0.006 | 0.010 | 0.008 |
| Bond angles (°) | 0.607 | 0.737 | 0.926 | 0.884 |
| Validation | | | | |
| MolProbity score | 1.81 | 2.02 | 2.35 | 2.39 |
| Clashscore | 6.3 | 7.30 | 10.65 | 11.91 |
| Poor rotamers (%) | 0.81 | 1.50 | 2.12 | 2.22 |
| Ramachandran plot | | | | |
| Favored (%) | 92.53 | 92.08 | 89.61 | 90.52 |
| Allowed (%) | 7.16 | 7.66 | 10.02 | 9.18 |
| Disallowed (%) | 0.31 | 0.27 | 0.38 | 0.30 |

**Supplementary Table 5. List of Ubiquitin peptides used in this study.**

| Abbreviation | Peptide sequence | Precursor m/z | Charge state | Product ions used for PRM | RT (min) |
|---|---|---|---|---|---|
| K6 linkage | M[Oxid]QIFVK[GG]TLTGK | 465.9270 | 3+ | y2+, y3+, y5+, y6+, y7+, y8+ | 39.1 |
| K6 linkage (heavy) | M[Oxid]QIFVK[GG]TLTGK | 468.2661 | | | |
| K11 linkage | TLTGK[GG]TITLEVEPSDTIENVK | 801.4269 | 3+ | y4+, y6+, y7+, y8+, y9+, y10+ | 42.5 |
| K11 linkage (heavy) | TLTGK[GG]TITLEVEPSDTIENVK | 803.4315 | | | |
| K27 linkage | TITLEVEPSDTIENVK[GG]AK | 701.0390 | 3+ | y6+, y7+, y8+, y9+, y10+ | 41.4 |
| K27 linkage (heavy) | TITLEVEPSDTIENVK[GG]AK | 703.0436 | | | |
| K29 linkage | AK[GG]IQDK | 408.7323 | 2+ | y2+, y3+, y4+, y5+ | 28.1 |
| K29 linkage (heavy) | AK[GG]IQDK | 412.2409 | | | |
| K33 linkage | IQDK[GG]EGIPPDQQR | 546.6129 | 3+ | y1+, y2+, y3+, y4+, y5+, y6+ | 31.8 |
| K33 linkage (heavy) | IQDK[GG]EGIPPDQQR | 548.6175 | | | |
| K48 linkage | LIFAGK[GG]QLEDGR | 487.6001 | 3+ | y2+, y3+, y4+, y5+, y6+, y8+ | 39.2 |
| K48 linkage (heavy) | LIFAGK[GG]QLEDGR | 489.9391 | | | |
| K63 linkage | TLSDYNIQK[GG]ESTLHLVLR | 748.7376 | 3+ | y5+, y6+, y7+, y8+, y9+ | 42.6 |
| K63 linkage (heavy) | TLSDYNIQK[GG]ESTLHLVLR | 751.0767 | | | |
| M1 linkage | GGM[Oxid]QIFVK | 448.2389 | 2+ | y2+, y3+, y4+, y5+, y6+, | 36.2 |
| M1 linkage (heavy) | GGM[Oxid]QIFVK | 451.2458 | | | |

**Supplementary Movie 1. Overview of the cryo-EM structures of human 26S proteasome in complex with the K11/K48-branched ubiquitin chain in different conformational states revealed by the study.**
